# Integrin-alpha-6+ Stem Cells (ISCs) are responsible for whole body regeneration in an invertebrate chordate

**DOI:** 10.1101/647578

**Authors:** Susannah H. Kassmer, Adam Langenbacher, Anthony W. De Tomaso

## Abstract

Colonial ascidians are the only chordates able to undergo whole body regeneration (WBR), during which entire new bodies can be regenerated from small fragments of blood vessels. Here, we show that during the early stages of WBR in *Botrylloides diegensis*, proliferation occurs only in small, blood-borne cells that express *integrin-alpha-6 (IA6), pou3* and *vasa*. Ablation of proliferating cells using Mitomycin C (MMC) blocks WBR in vascular fragments, but can be rescued by injection of cycling cells isolated from an untreated individual. Using prospective isolation and limit dilution analyes, we found that FACS-isolated IA6+ stem cells (ISCs) could rescue WBR in MMC treated vascular fragments, even when injecting only a single cell. Lineage tracing using EdU-labeling further revealed that donor-derived ISCs directly give rise to regenerating tissues. Inhibitors of either Notch or canonical Wnt signaling block WBR and reduce proliferation of ISCs, indicating that these two pathways regulate ISC activation.

## Introduction

The ability to regenerate missing structures is diverse and widespread among the metazoan phyla. Some invertebrate species within the phyla of Platyhelminthes, Cnidaria and Echinodermata, can regenerate whole bodies from small fragments of tissue, while some vertebrates such as amphibians and fish can regenerate ablated limbs and distal structures of some organs. In contrast, groups such as insects, birds and mammals have nearly lost the ability to regenerate (Li et al., 2015; Sanchez Alvarado, 2004; Sanchez Alvarado and Tsonis, 2006). Just like their general regeneration abilities, the cellular sources for regeneration are highly variable among regenerating species. In the Cnidarian *Hydractinia* and the Planarian *Schmidtea*, pluripotent stem cells are present in the adult and are responsible for regeneration, while in amphibians and zebrafish, lineage-committed progenitors arise from de-differentiation (Li et al., 2015; Sanchez Alvarado and Yamanaka).

In the majority of chordates, the ability to regenerate following a major injury is severely limited, usually resulting in scar formation. In contrast, a group of invertebrate chordate species; colonial ascidians of the genus *Botrylloides*, have been shown to regenerate whole bodies, including all tissues and organs, from small fragments of the vasculature. This process is called Whole Body Regeneration (WBR).

We are studying WBR in *Botrylloides diegensis*. Ascidians are the sister group of vertebrates, and begin life as a tadpole larvae with a typical chordate body plan. Following a free-swimming stage, the larvae settle and metamorphose into a sessile invertebrate adult, called a zooid. Zooids have a complex anatomy: they are filter feeders with siphons and a branchial basket, GI tract, central and peripheral nervous system, endocrine glands, a heart, and a circulatory system consisting of 8-12 blood cell types. In colonial ascidian species, zooids reproduce asexually, and the adult body plan consists of a colony of clonally derived individuals embedded within a gelatinous structure called a tunic (Figure 1A). The source of the new bodies is a specialized region of the body wall of the parental zooid called the peribranchial epithelium, and this process of asexual reproduction is called palleal budding (Kassmer et al., 2018). Asexual regeneration results in an ever-expanding number of zooids, embedded in the tunic, and linked by a common, extracorporeal vasculature (Figure 1A). This vascular bed extends beyond and encircles the zooids, and at the periphery of the colony terminates in structures called ampullae (Figure 1A).

**Figure 1:**
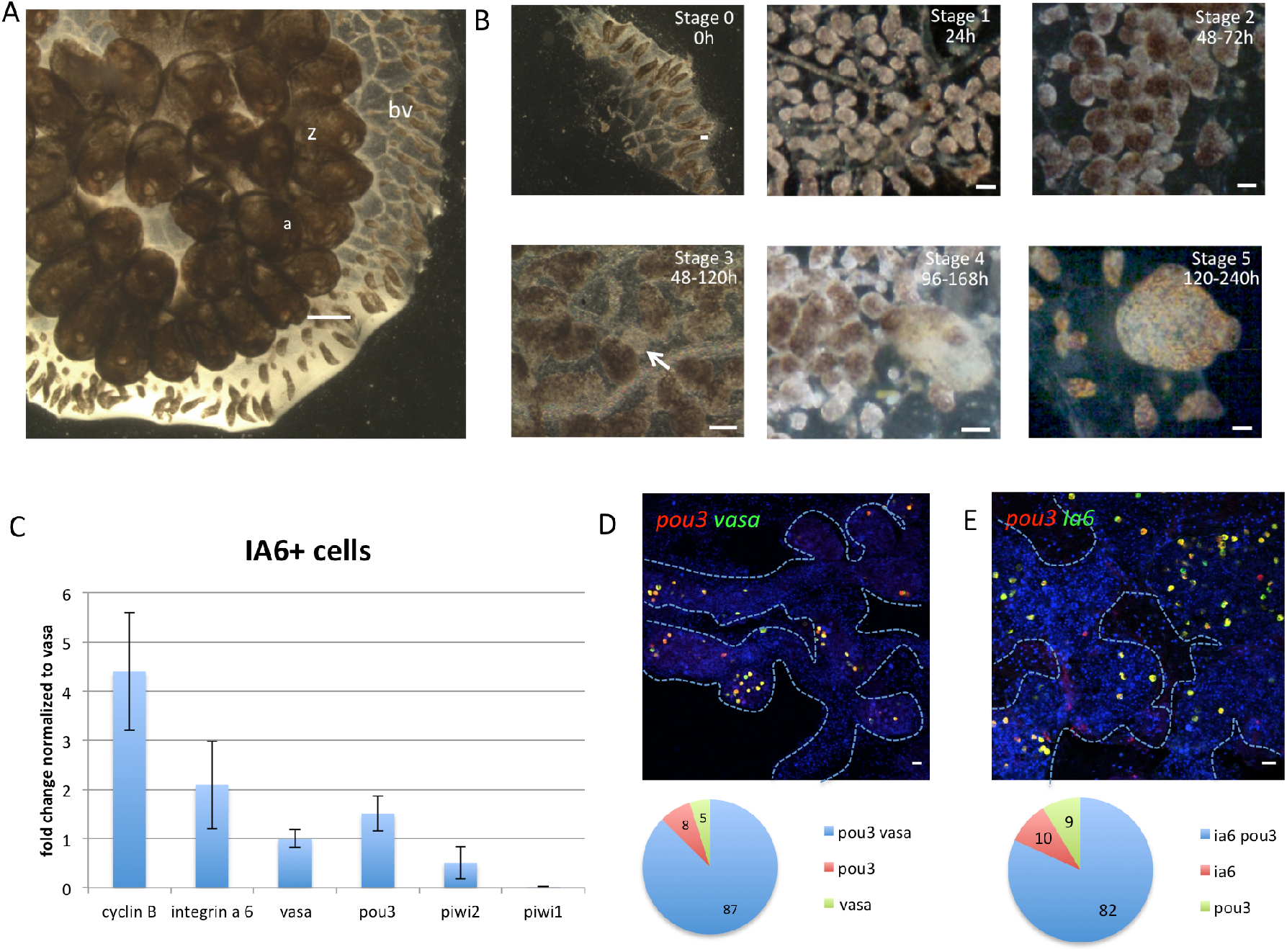
Stages of whole body regeneration, gene expression in IA6+ cells. A) *Botrylloides diegensis*, whole colony brightfield image, dorsal. Zooids (z) are embedded in a common, transparent tunic. Blood vessels (bv) extend throughout the tunic and are shared by the zooids of the colony. At the periphery of the colony, blood vessels form terminal sacs termed “ampullae” (a). Scale bar 1mm. B) Stage of whole body regeneration, brightfield images. Blood vessels are surgically separated from the colony (stage 0). During the first 24h, the blood vessel begins to remodel (stage 1). After 48h, blood vessels become condensed and highly pigmented (stage 2). Regeneration begins when a double vesicle is formed, consisting of two layers of epithelium (stage 3, white arrow). This double vesicle undergoes organogenesis (stage 4) and gives rise to a new, filter feeding body (stage 5). Scale bars 200um. C) qPCR analysis showing expression of *cyclin B, integrin-alpha-6, vasa, pou3, piwi2* and *piwi1* in IA6+ cells isolated by flow cytometry. Data are expressed as averages of fold changes normalized to *vasa*, n=4. Error bars show standard deviation. D) FISH showing coexpression of *pou3* (red) and *vasa* (green) in stage 0. DNA was stained with Hoechst (blue). Blue dashed lines outline blood vessel boundaries. Scale bar 20μm. Single positive and double positive cells were counted using the cell counter feature in FIJI, and for each stage, 4 images from 4 independent samples were counted. Pie graph shows averages of percentages of *pou3/vasa*-double-positive cells as well as *pou3* and *vasa* single positive cells. E) FISH showing co-expression of *pou3* (red) and *ia6* (green) in stage 0. DNA was stained with Hoechst (blue). Blue dashed lines outline blood vessel boundaries. Scale bar 20μm. Single positive and double positive cells were counted using the cell counter feature in FIJI, and for each stage, 4 images from 4 independent samples were counted. Pie graph shows averages of percentages of *pou3/ia6*-double-positive cells as well as *pou3* and *ia6* single positive cells.

In addition to palleal budding, which occurs normally, *B. diegensis* can also use the alternative WBR pathway to produce a zooid following injury. WBR occurs in the vasculature, and is initiated by surgical removal of all zooids, or separation of small vascular fragments from the rest of the colony (Figure 1B). WBR progresses through distinct visual stages. For the first 24 hrs following surgical ablation, the blood vessels undergo regression and remodeling (stage 1, Figure 1B). During this time, blood flow, usually powered by the hearts of each zooid, continues and is driven by contractions of the remaining ampullae (Blanchoud et al., 2017). During the next 48 hrs, the blood vessels continue to remodel and form a dense, contracted network (stage 2, Figure 1B). Within this dense, highly pigmented vascular tissue, an opaque mass of non-pigmented cells becomes apparent; creating a clear area that is the presumptive site of bud development (stage 3, arrow in Figure 1B). The mass of cells next forms into a hollow, blastula-like epithelial sphere. The vascular epithelium then wraps itself around this sphere, leading to the formation of a distinct visible double vesicle. Over the next 48h, the inner vesicle increases in size (Stage 4, Figure 1B), while undergoing a series of invaginations and evaginations that lead to organogenesis and the eventual regeneration of a zooid. WBR is defined as complete when the new zooid is actively filter-feeding, and occurs within a range of 7-10 days (stage 5; Figure 1B). The zooid immediately commences normal palleal budding, and the colony regrows.

In the present study, we aimed to assess the cellular origins of WBR in *Botrylloides diegensis*, specifically the role of circulatory cells in this process. WBR has been studied in three different species of botryllid ascidians, and in all cases a population of cells with an undifferentiated appearance, termed hemoblasts, have been suggested to initiate this regenerative process (Brown et al., 2009a). In *Botrylloides violaceus*, 15-20 small hemoblasts that express Piwi protein have been shown to aggregate under the epidermis of a blood vessel during early WBR (Oka, 1959). These cells can be detected during the early vesicle stage and occasionally within the epithelium of a vesicle, suggesting that they play a role in regeneration (Brown et al., 2009a). In a related species, *Botrylloides leachii*, Blanchoud et al. showed an increase in the population of hemoblasts very early after injury during WBR (Blanchoud et al., 2017). These results had suggested that blood borne stem cells might play a role in WBR in *Botrylloides*, but to date, it has never been directly tested whether such cells give rise to regenerating tissues.

In several species of colonial ascidians, the blood contains self-renewing, lineage restricted germline stem cells that migrate to newly developing buds during repeated cycles of asexual reproduction, where they give rise to eggs and testes (Brown et al., 2009b; Carpenter et al., 2011; Kawamura and Sunanaga; Laird et al., 2005; Rodriguez et al., 2016; Sunanaga et al., 2010; Sunanaga et al., 2006). In *Botryllus schlosseri*, these germline stem cells can be enriched by flow cytometry using Integrin-alpha-6 (IA6) as a marker (Kassmer SH, 2015) and express *piwi* as well as other genes associated with germ cells, such as *vasa*, and *pumilio (Brown et al., 2009b; Langenbacher and De Tomaso, 2016; Sunanaga et al., 2006)*. Since *vasa* and *piwi* are part of the germline multipotency program (Juliano et al., 2010), and IA6 is a biomarker for various kinds of mammalian stem cells, including embryonic stem cells and primordial stem cells (Krebsbach and Villa-Diaz, 2017), we hypothesized that IA6 might likewise be expressed on blood borne stem cells that are involved in WBR in *Botrylloides*.

In order to isolate the cells responsible for WBR, we developed a rescue assay and used a prospective isolation strategy to identify the cells that give rise to regenerating tissues. We show that during the very early stages of WBR in *B. diegensis*, cell proliferation occurs only in blood-borne cells that express *integrin-alpha-6, pou3, vasa and piwi*. IA6+ cells are required for WBR and lineage tracing using EdU labeling reveals that they directly give rise to regenerating tissues. Finally, we show that proliferation of IA6+ cells during WBR is regulated by Notch and Wnt signaling.

## Results

### The majority of proliferating cells during early stages of WBR express Integrin-alpha-6, vasa and pou3

To assess whether Integrin-alpha-6 enriches for cells expressing stem cell-associated genes in *B. diegensis*, we isolated IA6+ cells from the blood of control genotypes by flow cytometry and quantified the expression of germline pluripotency genes such as *vasa*, *piwi1, piwi2* and *pou3*. For the latter, we hypothesize that the octamer binding transcription factor Pou3 plays a role in stem cell pluripotency in ascidians, similar to Oct4 in mammalian pluripotent cells. Oct4 is a member of the Pou class 5 gene family; a vertebrate specific family of pou genes (Sanchez Alvarado and Yamanaka; Shi and Jin, 2010). It is likely that *pou5* inherited this function from an ancestral *pou* paralog (Gold et al., 2014), and in the cnidarian *Hydractinia echinata, pou3* plays a role in stem cell pluripotency (Millane et al., 2011). We cloned *pou3* from *B. diegensis* and *B. schlosseri* and constructed a phylogenetic tree that confirms the close relationship of *pou3* to *pou5* (Figure S1). *Pou3* is expressed in the developing germline of palleal buds in *B. diegensis* colonies (Figure S1), and *vasa* and *pou3* are highly expressed in IA6+ cells isolated from the blood (Figure 1C). Previously, only one *piwi* gene had been reported in *B. leachii (Rinkevich et al., 2010)*, but we found that, like most animals, *Botrylloides diegensis* has two *piwi* genes (Juliano et al., 2011; Seto et al., 2007). Only *piwi2* is expressed in IA6+ cells, at lower levels than *vasa* (Figure 1C). Using double-labeled FISH, we confirmed that 81% of either *ia6*+ or *pou3*+ cells co-express both genes, while 9 and 10% express only *pou3* or *ia6*, respectively (Figure 1E). The overlap between *vasa* and *pou3* is 87% (Figure 1D).

To assess whether *ia6*+ *pou3*+ cells are involved in responding to injury, we analyzed the expression of these two genes in proliferating cells during the early time points after surgical separation of blood vessels from the colony. We used *Histone 3* mRNA expression as a marker of proliferating cells, as it is upregulated in S-phase of the cell cycle in plants and animals, and has been used as a marker for cell proliferation in in situ hybridization previously (Arakura et al., 2001; Gown et al., 1996; Langenbacher et al., 2015; Meshi et al., 2000; Osley, 1991). Analyzing expression of *histone 3 (h3)* together with *integrin-alpha-6* by double fluorescent in situ hybridization (FISH), we found that 90% of all *ia6*+ cells proliferate in the blood of a healthy colony (stage 0, Figure 2 A and B). After separation of blood vessels form the colony, the overlap between *h3* and *ia6* stays high during stages 1 and 2 (86% and 92%, respectively, (Figure 2A and Figure S1E), and the total number of proliferating *ia6*+*h3*+ cells increases dramatically (Figure 2A, compare stage 0 and 2). Furthermore, *ia6*+ cells make up the majority of proliferating cells during the early stages of WBR, as the number of *h3* single positive cells is consistently much lower than that of *ia6*+*h3*+ double positives in stages 0, 1 and 2 (Figure 2B). During stages 1 and 2, some *ia6*+ cells aggregate (white arrows in Figure 2, Stage 2), a structure that we refer to as regeneration foci. Cells in the aggregates continue to proliferate, and the structure increases in size (Figure 2, Stage 3, yellow arrows). The aggregate next begins to form the epithelial sphere (Figure 2, Stage 3, red arrows), which will eventually form the zooid. As the aggregates increase in size, a concurrent drop in IA6 expression can be seen (Figure 2, Stage 3, yellow arrows) and by early vesicle phase there is almost no IA6 expression (Figure 2, Stage 3, red arrows). This is likely a sign of differentiation of these cells. In contrast, IA6+ cells that are not present in developing epithelial spheres continue to proliferate (Figure 2A,B).

**Figure 2:**
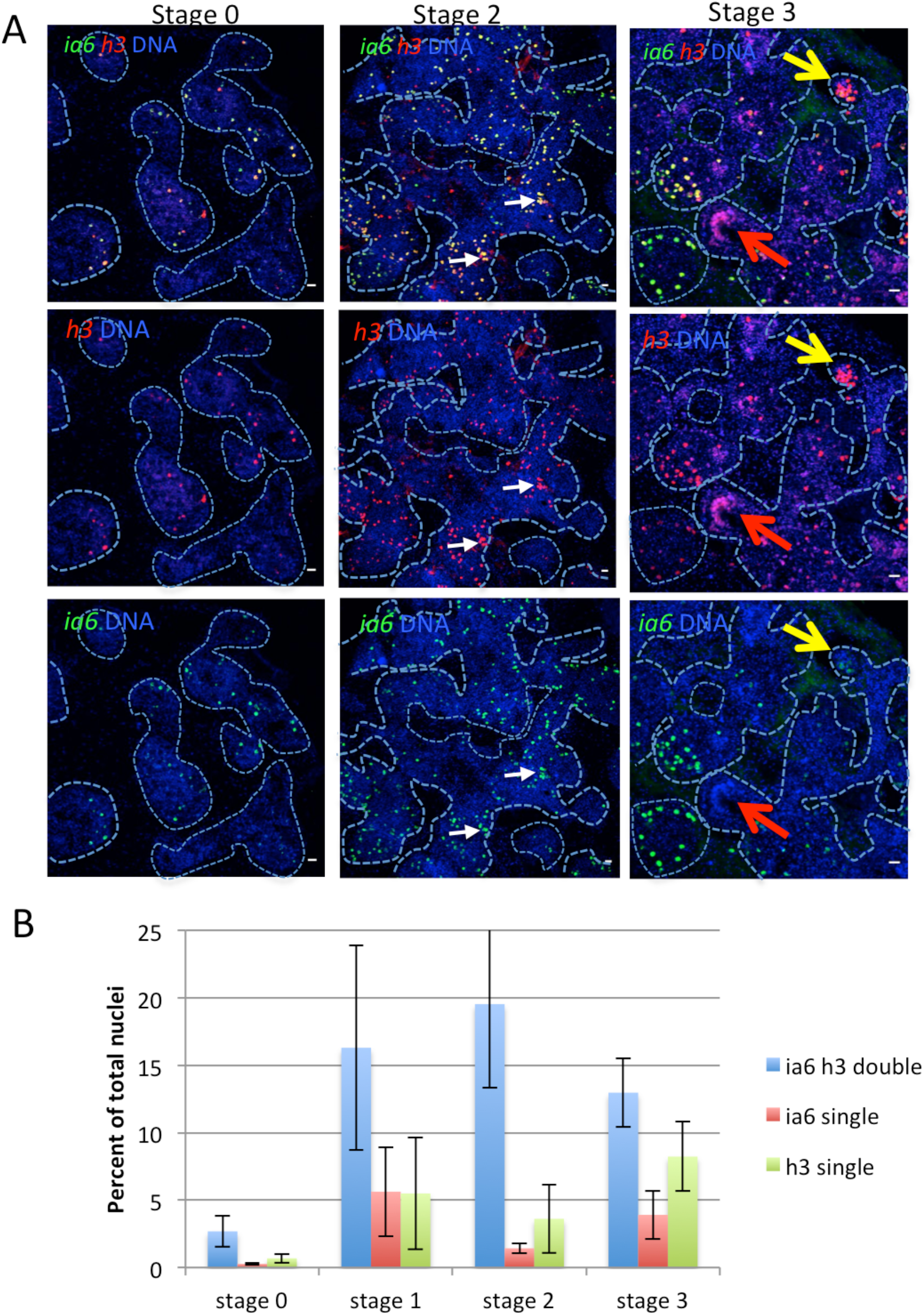
Whole body regeneration is associated with proliferation of blood borne *integrin-alpha-6*+ cells. A: Fluorescent in situ hybridization (FISH) showing expression of *integrin-alpha-6 (ia6*, green) and *histone 3* (*h3*, red) during stages 0, 2 and 3 of WBR. White arrows indicate *ia6*+ cells beginning to cluster during Stage 2. Yellow and red arrows in Stage 3 show different stages of WBR. As clusters increase in size, they begin to lose *ia6* expression (yellow arrows). As cell clusters differentiate to form the blastula-like structure, *ia6* expression is not detected (red arrows). DNA was stained with Hoechst (blue) in all panels. Blue dashed lines outline blood vessel boundaries. Scale bars 20μm. B) Single positive (*ia6* or *h3*) and double positive cells were counted using the cell counter feature in FIJI, and for each stage, 4 images from 4 independent samples were counted. Graph shows percentages of *ia6/h3* double positive and *ia6*− or *h3*-single positive cells among all Hoechst-positive nuclei. Error bars show standard deviation.

Analyzing expression of *h3* and *pou3* by double FISH, we found that proliferating cells in the blood of *Botrylloides* express *pou3* in uninjured steady state and during the early stages of regeneration (Figure S2). 91% of all *pou3*+ cells proliferate in steady state, and this percentage stays high during stages 1 and 2. By stage 3, 64% of *pou3*+ cells become quiescent, and cell proliferation is predominant (71%) in *pou3*− regenerating double vesicles (yellow arrows in Figure S2, stage 3) and blood cells (Figure S2, stage 3). Together, these results show that *ia6+/pou3*+ cells proliferate in steady state and make up the majority of proliferating cells during the early stages of WBR. Based on these findings, we hypothesize that *ia6*+*pou3*+ proliferating cells are stem cells responsible for WBR.

### IA6+ cells are required for regeneration

To functionally assess whether proliferating IA6+ cells are involved in whole body regeneration, we ablated proliferating cells using the drug Mitomycin C (MMC). Vascular tissue was soaked in MMC immediately after surgery and treated for 24h until stage 1, when overlap between IA6 and the proliferation marker histone 3 is 93% (Figure 2). In MMC treated vessels at 5 days post treatment, no *pou3*+ or *histone3*+ cells remain (Figure S3A). Elimination of proliferating cells with Mitomycin C at stage 0-1 results in subsequent loss of regenerative capacity (Figure 3A). Normally, regeneration is complete after 11-14 days (Figure 1), but MMC treated samples do not reach stage 4 or stage 5 even after 20 days (Figure 3A), and most MMC treated blood vessels appear to be permanently arrested in stage 2 (Figure 3A). However, regeneration can be rescued in MMC treated vasculature by injection of 2000 total blood cells isolated from an untreated, healthy individual, demonstrating that WBR depends on blood borne cells (Figure 3B). To identify the population of cells responsible for this rescue, we injected different populations of cells isolated by flow cytometry and assessed rescue of WBR (Figure 3C). We initially compared cells in the G2/M phase of the cell cycle versus those in G0, and found that 50 cycling cells could rescue regeneration, while 50 G0 cells could not. We next compared IA6+ and IA6- populations. While 50 IA6- cells are not able to rescue WBR, injection of 1000 IA6- cells achieved a 57% rescue efficiency (Figure 3C). This could be due to cell-sorting impurities or it could suggest that a rare population of IA6- stem cells exists (about 1/2000), which is supported by the finding that almost 20% of *pou3*+ cells are *ia6-* (Figure 1E). In contrast, 90% rescue is achieved by injecting only 50 IA6+ cells (Figure 3C). We next carried out limiting dilution analyses of the IA6+ population, and found that injection of a single IA6+ cell isolated from the blood of a healthy animal can rescue whole body regeneration in 20% of MMC treated samples (Figure 3C). Using ELDA analysis software (Hu and Smyth, 2009), we calculated the estimated frequency of IA6+ cells capable of rescuing WBR to be about 1 in 7.55 (upper estimate 4.63, lower estimate 12.3). These results show that IA6+ cells can independently rescue WBR.

**Figure 3:**
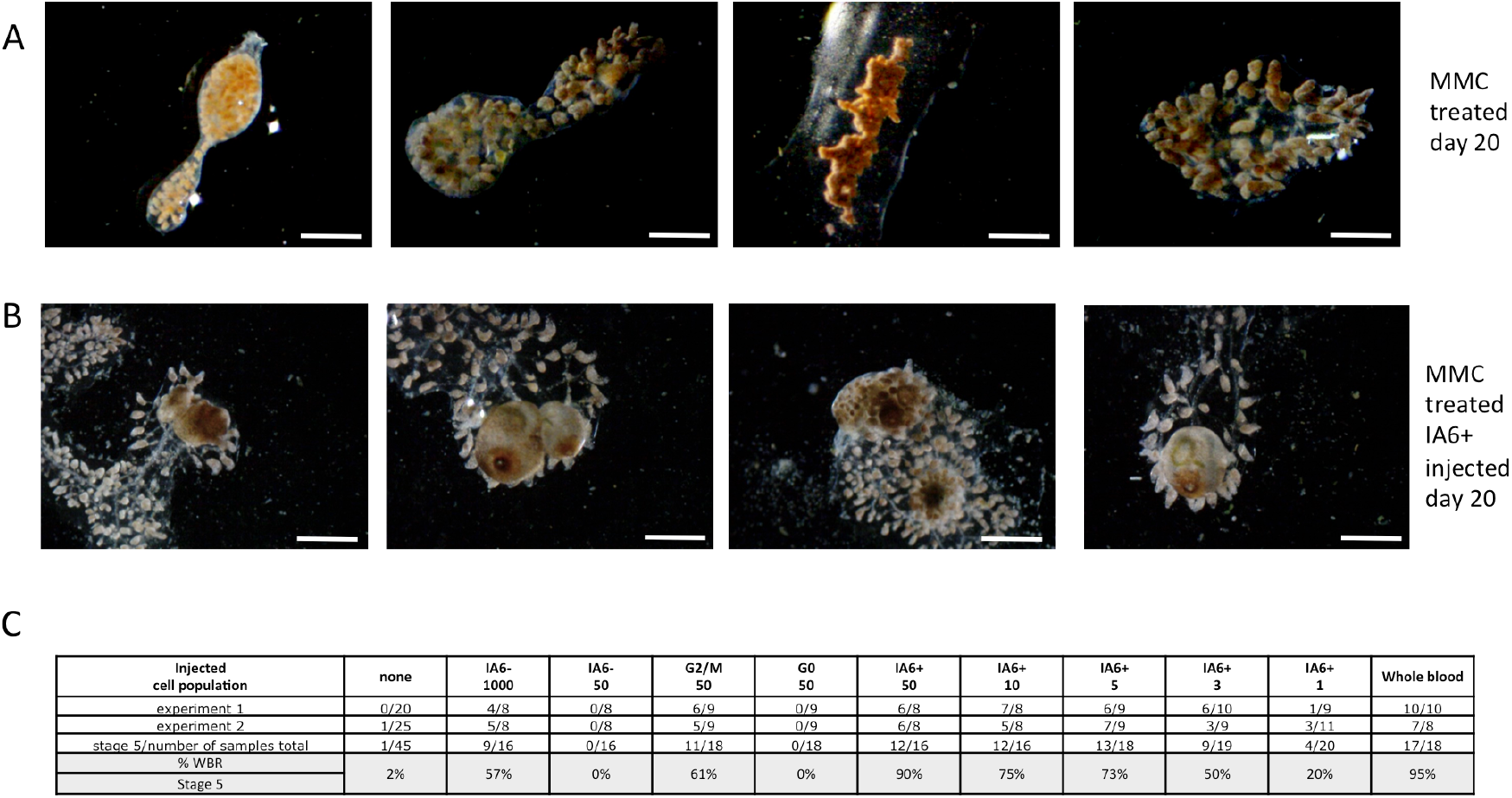
IA6+ cells are required for whole body regeneration. A: Mitomyinc C (MMC) treatment prevents regeneration. Brightfield images of vessel fragments 20 days post MMC treatment. Vessel fragments were treated with MMC for 16h immediately after surgery. After 16h, MMC was removed, and the samples followed for 20 days. Scale Bar 1mm B: Injection of IA6+ cells 24h after MMC treatment rescues WBR. Brightfield images of MMC treated vessel fragments 20 days post injection of IA6+ cells. Scale Bar 1mm C: Table showing rates (percentages) of WBR (reaching stage 5) 20 days after injection of different cell populations. For each condition, 16-20 samples were injected in two independent experiments.

### IA6+ Stem Cells (ISCs) give rise to regenerating tissues in developing bodies

Since ISCs appear to be functionally required for WBR, we wanted to assess whether progeny from IA6+ cells is incorporated into regenerating tissues. We used EdU to label IA6+ cells in steady state in an intact animal and subsequently track the presence of their progeny in regenerating tissues by transplanting them into MMC treated vessel fragments. The modified thymidine analogue EdU is incorporated into newly synthesized DNA and detected in fixed tissues using a fluorescent dye. We ensured that EdU could be detected at stage 3 of WBR in vessels injected with EdU at stage 0 (Figure 4, middle), while uninjected samples showed no signal (Figure 4, left). IA6+ cells were isolated from donor animals that had been injected with EdU the day before to allow EdU incorporation into the DNA of cycling cells. EdU-labeled IA6+ cells were injected into MMC treated recipients as in the rescue experiments described above. When recipients reached stages 2-4 of WBR, the regenerating tissues were fixed and stained for EdU (Figure 4, right). By stage 2, EdU+ cells were found in cell aggregates of regeneration foci (Figure 4, right) as well as in circulation (Figure S3). Examples of three independent experiments are shown in Figure S3, At Stage 4, EdU+ cells could be identified in the epithelia of the regenerating body that are undergoing morphogenesis (Figure S3B, white arrows). In one case, we also saw EdU+ cells either part of or directly underneath the outer epithelium of the regenerating body (Figure S3B, green arrow). These results show that IA6+ cells are the source of regenerating tissues in newly developing bodies.

**Figure 4:**
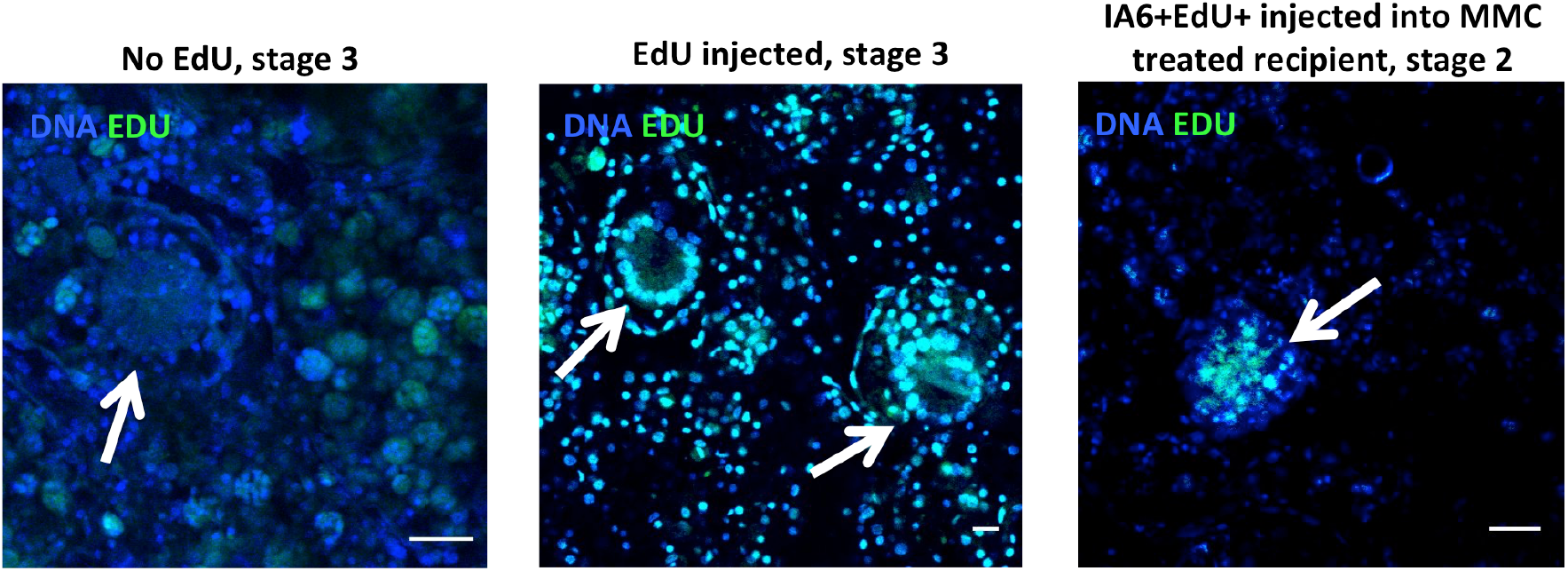
IA6+ cells give rise to regenerating tissues. Overlay of nuclear staining (Hoechst, blue) and Edu (green). Left: In vessel sections from stage 3 samples not injected with EdU, no green EdU signal is detected. Middle: In stage 3 samples that have been injected with EdU at stage 0, EdU-positive cells are present in the blood as well as in regenerating double vesicles (white arrows). Right: IA6+ cells were isolated from Edu-treated animals and injected into MMC treated vessel fragments. Edu-positive cells give rise to regeneration foci at late stage 2 (white arrows). Scale bars 20μm. Images are representative of 10 samples from two independent experiments.

### Inhibition of Notch or Wnt signaling blocks regeneration and proliferation of ISCs

We next aimed to analyze the expression of components of the Notch and Wnt signaling pathways in proliferating cells from the blood during the early time points of WBR. These pathways are known to play roles in regulating stem cell activity, proliferation of blastema cells and cellular differentiation in other organisms (Grotek et al., 2013; Hamada et al., 2015; Kawakami et al., 2006; Perdigoto and Bardin, 2013; Reya and Clevers, 2005; Tal et al., 2010; Vogg et al., 2016; Wehner et al., 2014) and are upregulated during WBR in *Botrylloides* (Zondag et al., 2016). We isolated cycling G2/M cells by flow cytometry from stage 0 as well as at 24h and 48h after injury of vessel fragments. We specifically isolated G2/M cells instead of IA6+ cells in order to be able to analyze gene expression in all proliferating blood cells, even those that may have begun to down-regulate IA6 as they differentiate (Figure 2). As expected, G2/M cells isolated from the blood of healthy, unmanipulated animals (0h), express high levels of *ia6*, mitosis-specific *cyclin b*, and *pou3* (Figure 5A). *Notch 1, notch 2* and the downstream gene *hes1*, as well as the Wnt pathway components *frizzled5/8, disheveled* and *beta catenin*, are also expressed, indicating active Notch and Wnt signaling in proliferating circuating cells from control individuals (Figure 5B). 24h after injury, *notch2, frizzled 5/8* are upregulated in G2/M cycling cells compared to 0h. 48h after injury, *notch2, pou3* and *frizzled 5/8* are even more highly upregulated. *Notch1* is downregulated at 24h and 48h (Figure 5C). *Piwi2* is highly upregulated in cycling G2/M cells at 48h together with *vasa* (Figure S4). These results show that the pluripotency/stem cell related genes *pou3, ia6, piwi2* and *vasa* are highly expressed in cells that proliferate during the early stages of WBR. Double FISH for *notch1* and *h3* shows that *notch1*+ cells proliferate during stage 0 and stage 1, but fewer of these cells are present in stage 2 (Figure 6A). However, the number of *notch2*+ cells increases by stage 2 (Figure 6A). These results suggest that *notch1* is primarily responsible for regulating proliferation of ISCs during steady state and at early time points post injury. In Stage 2, notch1-positive cells are more quiescent and fewer in number. At the same time, notch2 is expressed in a higher proportion of proliferating cells. This could be related to a role of notch2 in regulating proliferation of differentiating progenitors, while notch1 is associated with stem cell maintenance.

**Figure 5:**
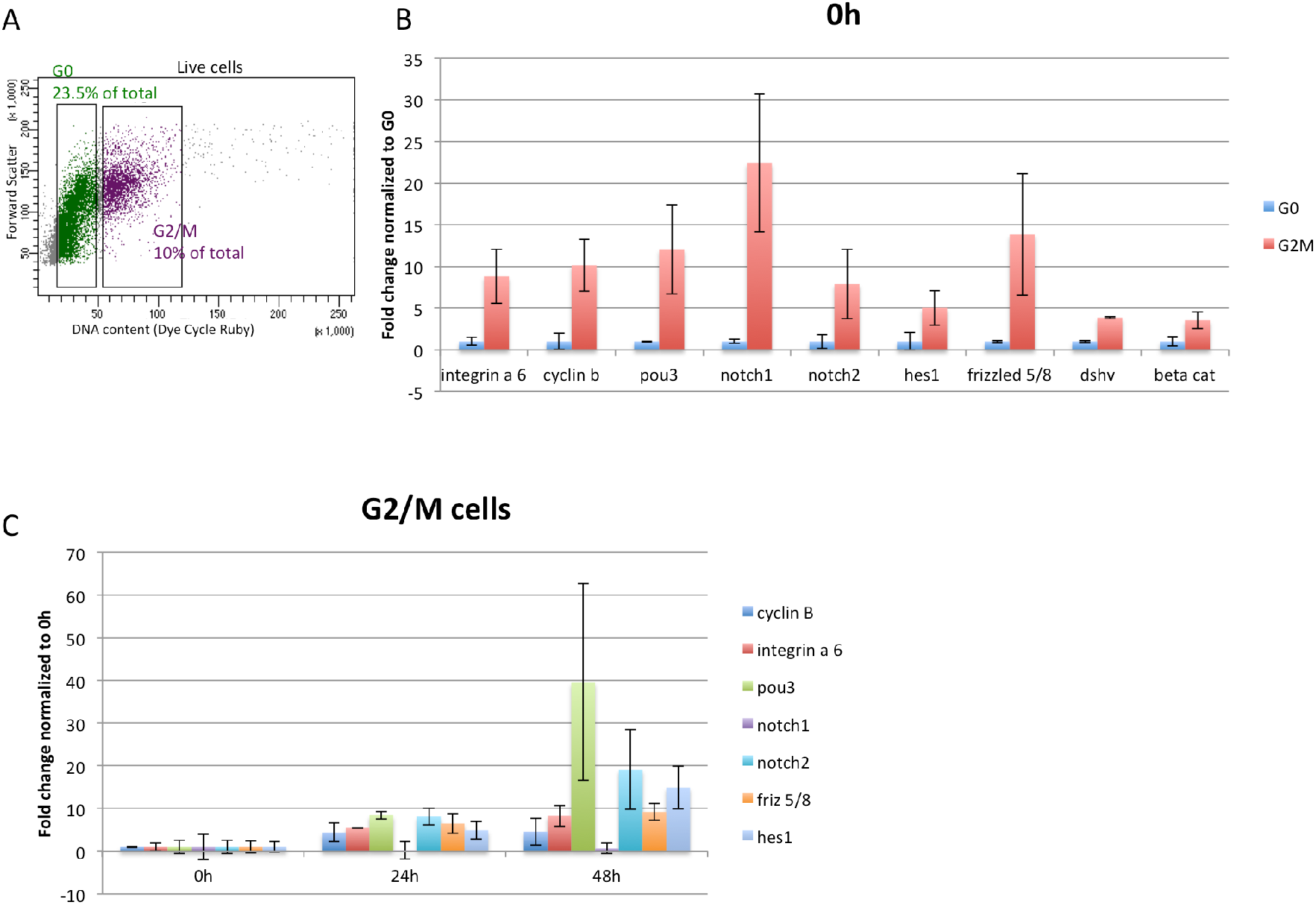
Gene expression in cycling blood borne cells in steady state and during WBR. A) Isolation of cycling G2/M cells by flow cytometry. Cells were stained with the live DNA stain Dye Cycle Ruby. Linear analysis of fluorescence intensity plotted against forward scatter (cell size) shows a clear separation of G2/M cells with 4n DNA content and G0/G1 cells with 2n DNA content. G2/M cells comprise 10% of live cells. B) qPCR analysis showing expression of, *integrin-alpha-6, cyclin b, pou3, notch1, notch2, hes1, frizzled 5/8, disheveled (dshv)* and *beta catenin* in G2/M cells in steady state (0h). Data are expressed as averages of fold changes normalized to G0 cells, n=3. C) qPCR analysis showing expression of *cyclin b, integrin-alpha-6, pou3, notch1, notch2*, and *frizzled 5/8* in G2/M cells at 24h and 48h post-injury, Data are expressed as averages of fold changes normalized to 0h (stage 0), n=4. Error bars represent standard deviation.

To assess whether Notch or Wnt signaling are required for WBR, vessel fragments were allowed to regenerate in the presence of inhibitors of either Notch (DAPT) or Wnt signaling (Endo-IWR). Inhibition of either Notch or canonical Wnt signaling blocked regeneration in a dose dependent manner (Figure 6B). In either 2uM of the Notch-signaling inhibitor DAPT or 1uM of the canonical Wnt signaling inhibitor Endo-IWR, the vessel tissue underwent remodeling up to stage 2 in both treatments, but never progressed beyond stage 2 while kept in drug (for up to 4 weeks), and appeared otherwise healthy and alive. Upon removal of the inhibitors (after 96h of treatment), regeneration progressed normally (14/15 samples reached stage 5 for IWR and 15/16 for DAPT, Figure 6B), indicating that regeneration is only halted, but that all cell types required for WBR are still viable and fully functional. As controls, we used Exo-IWR which is a 25-fold less active against the Wnt/β-catenin pathway, and the gamma-secretase modulator E2012, which does not affect the Notch specific gamma-secretase. Both control drugs Exo-IWR (1uM) and E2012 (1uM) did not affect regeneration rates (Figure 6B).

**Figure 6:**
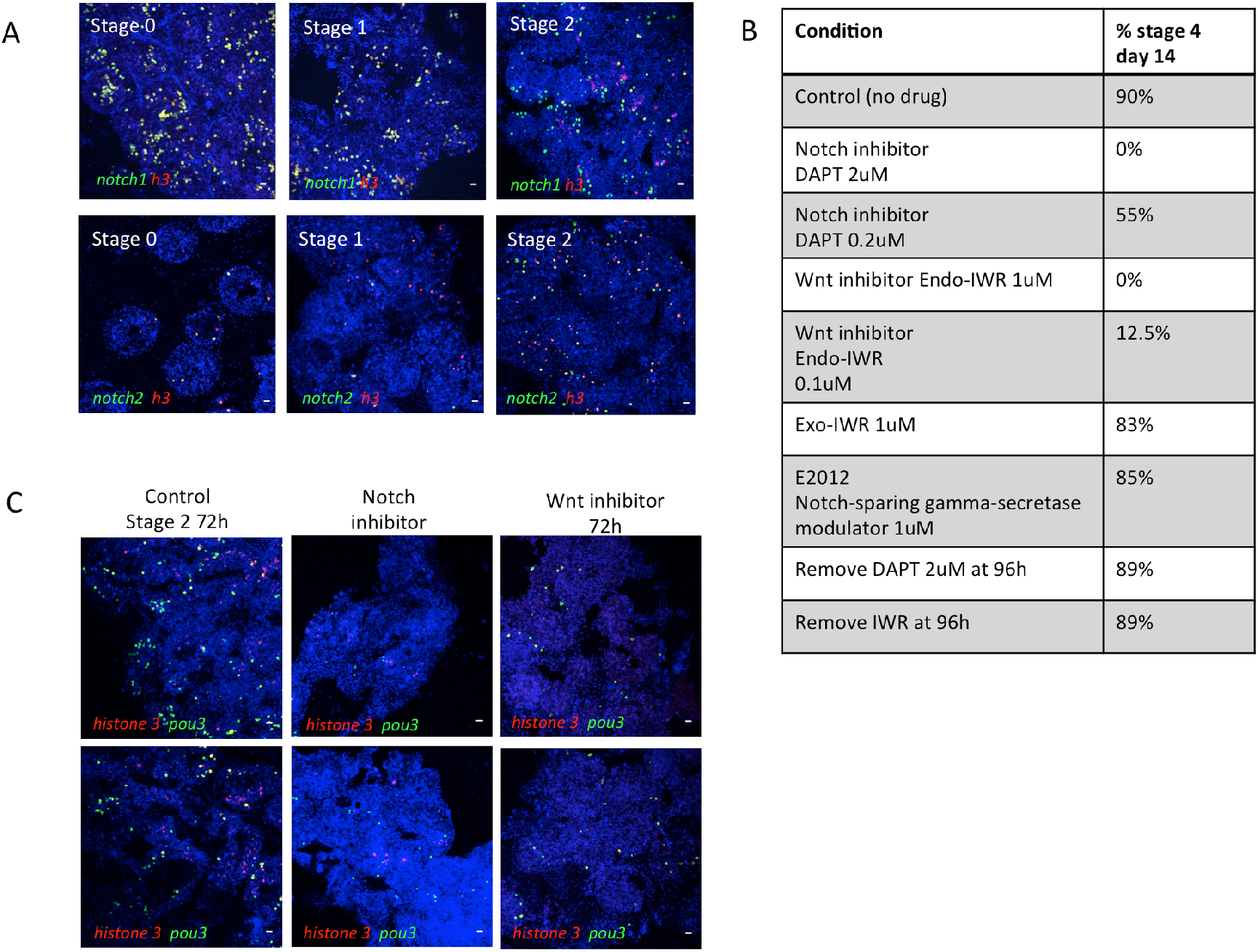
Inhibition of Notch or Wnt signaling prevents WBR and blocks proliferation of pou3+ cells. A: top panel: FISH for *notch1* (green) and *h3* (red) during steady state (stage 0) and during early stages of WBR. Bottom panel: FISH for *notch2* (green) and *h3* (red) during steady state (stage 0) and during early stages of WBR. Scale bar 20μm B: table showing the percentages of vessel fragments reaching stage 4 at day 14 post injury during treatment with different doses of inhibitors of canonical wnt signaling (Endo-IWR), Notch signaling (DAPT) or control drugs Exo-IWR (1μM) and the Notch-sparing gamma-secretase modulator E2012 (1μM). The number of colonies that reached stage 4 were counted at 14 days post injury. Percentages represent the number of fragments reaching stage 4, with n= 14-18 fragments for each condition. Controls received vehicle only. C) FISH for *pou3* (green) and *h3* (red) showing a reduction in proliferating *pou3*+ cells 72h after surgery when treated with an inhibitor of either Notch or Wnt signaling. Scale bars 20μm.

We next assessed for the presence of proliferating cells in drug treated vessel fragments using double FISH for *h3* and *pou3*. Following 72h of treatment, fewer *pou3*-positive cells are present in drug treated vessels compared to controls (Figure 6C), and regeneration foci do not form. Of the few pou3+ cells that remain, very few (Notch inhibitor) to none (Wnt inhibitor) are proliferating. We conclude that inhibition of signaling downstream of Wnt and Notch receptors blocks ISCs from entering the cell cycle, and pushes them towards quiescence. Because ISCs remain quiescent, they are unable to form regeneration foci. Inhibition of Notch or Wnt signaling likely also affects differentiation of ISCs, as there are much fewer proliferating pou3- cells compared to controls. Yet, upon removal of the inhibitors, ISCs are able to exit quiescence and to resume proliferation and differentiation normally. In addition, whatever cell type is producing the Wnt and Notch ligands is still capable of doing so when the inhibitors are removed. Together, these results show that WBR depends on both Wnt and Notch signaling, and that both signaling pathways control the response of ISCs to injury and affect their proliferation and differentiation.

## Discussion

Here, we provide definitive functional evidence that blood-borne stem cells are responsible for injury induced whole body regeneration (WBR) and give rise to regenerating tissues in an invertebrate chordate. These cells belong to the undifferentiated “hemoblast” population that has been described historically in the blood of colonial ascidians (Brown and Swalla, 2012) (Kawamura and Sunanaga) and has been implicated in whole body regeneration in previous descriptive studies (Brown et al., 2009a; Oka, 1959).

At the level of a single cell, we have found that IA6+ Stem Cells (ISCs) can give rise to regenerating tissues during WBR, and that WBR cannot proceed when ISCs are ablated. We show that ISCs are constantly dividing in steady state and express genes associated with pluripotency, such as *pou3*. ISCs also express germ plasm components such as *vasa* and *piwi*, yet we have shown that they directly give rise to somatic tissues during WBR. So far, we have developed two main hypotheses about the nature of these cells.

The first hypothesis is that *Botrylloides* maintains a pluripotent stem cell used for WBR. In several other invertebrate species, germ plasm components such as *vasa, nanos* and *piwi* are expressed in cells which have somatic potential and have therefore been defined as germline multipotency (GMP) genes (Juliano et al., 2010). These GMP expressing cells are called Primordial Stem Cells and are part of the germline and can be involved in asexual reproduction in some animals (Solana, 2013). Examples of germ plasm containing cells that also give rise to somatic tissues include small micromeres from sea urchins, neoblasts in planarians, i-cells in hydrozoan cnidarians and archeocytes in sponges (Juliano et al., 2014; Millane et al., 2011; Reddien, 2018; Reddien and Sanchez Alvarado, 2004; Voronina et al., 2008; Wagner et al., 2011; Yajima and Wessel, 2011). One hypothesis is that that at least a subset of ISC’s might be similar to these types of primordial stem cells, and maintain somatic potential. Both the low frequency of rescue of single IA6 cell transplants, and the rare IA6- cells that also can rescue suggest that there is underlying heterogeneity in these cell populations.

Another hypothesis is that a normally lineage restricted germline progenitor is triggered to proliferate and differentiate into somatic tissues under certain conditions, such as loss of all zooids. This process might be somewhat similar to teratoma formation in mammals. Teratomas are germ cell tumors that contain an assortment of tissue structures from all three germ layers: endodermal, mesodermal, and ectodermal. These tumors are derived from primordial germ cells that migrate to ectopic sites during embryogenesis and differentiate into somatic cell types. In *Botrylloides*, the mechanisms that normally restrict circulatory germline progenitors to germ cell fate might be temporarily lifted following injury and subsequent whole body regeneration. *Vasa*− positive germline stem cells have been described in the blood of several botryllid ascidians and persist throughout adult life (Brown and Swalla, 2007; Kawamura and Sunanaga, 2011; Sunanaga et al., 2007). In *B. schlosseri*, we have shown that these cells are lineage restricted and give rise to new gonads during repeated rounds of asexual reproduction (Brown et al., 2009b; Carpenter et al.; Laird et al., 2005), and can be isolated based on expression of IA6 (Kassmer et al., 2015). While we have not functionally demonstrated that IA6+ cells in *B. diegensis* are lineage restricted (B. diegensis is not constantly fertile in lab-reared conditions), they express equivalent pluripotency genes. Thus, upon separation of vessels from bodies in *Botrylloides*, the pluripotency of IA6+ cells may be utilized for somatic regeneration. In contrast, all somatic tissues that are formed during asexual reproduction (palleal budding) in healthy animals are derived from the peribranchial epithelium of the parental zooids and not from blood borne cells (Berrill, 1947; Brown and Swalla, 2012; Carpenter et al., 2011; J., 1951; Oka, 1959).

In a large-scale gene expression study, Zondag et al. showed that Wnt and Notch signaling components are upregulated during early stages of regeneration in *B. leachii (Zondag et al., 2016)*. Here, we show that these pathways are required for proliferation and possibly maintenance of ISCs. Both of these pathways have been shown to play roles in regulating regeneration and stem cell proliferation in several other regenerating species (Sanchez Alvarado and Tsonis, 2006), and Wnt signaling is required for blastema formation (Kawakami et al., 2006) and stem cell maintenance (Reya and Clevers, 2005). Notch regulates blastema cell proliferation during zebrafish fin regeneration, and it mediates cell fate decisions such as proliferation and differentiation in many stem cell types (Perdigoto and Bardin, 2013). Notch also regulates blastema formation during distal regeneration in the solitary ascidian *Ciona* (Hamada et al., 2015). Future studies will investigate which Notch and Wnt ligands are expressed by the regeneration niche in *Botrylloides*, and how and when these signaling pathways act on ISCs to regulate proliferation and differentiation.

In summary, we have identified a stem cell population responsible for whole body regeneration in a chordate species, *Botrylloides diegensis*, and developed the toolsets necessary for future detailed molecular analysis of the stem cells involved in WBR and the pathways that regulate their function.

## Methods

### Animals

*Botrylloides diegensis* colonies used in this study were collected in Santa Barbara, CA and allowed to attach to glass slides. Colonies were maintained in continuously flowing seawater at 19-23°C and cleaned with soft brushes every 14 days.

### Whole body regeneration

Under a stereo dissection microscope, blood vessel fragments were surgically separated from the rest of the colony using razor blades. Regenerating fragments were kept attached to slides in flowing filtered seawater at 19-23°C.

### Integrin-alpha 6 and cell cycle flow cytometry

Genetically identical, stage matched samples were pooled, and blood was isolated by cutting blood vessels with a razor blade and gentle squeezing with the smooth side of a syringe plunger. Blood was diluted with filtered seawater and passed through 70 μm and 40 μm cell strainers. Anti-Human/Mouse-CD49f–eFluor450 (Ebioscience, San Diego, CA, USA, cloneGoH3) was added at a dilution of 1/50 and incubated on ice for 30 min and washed with filtered seawater. For cell cycle sorting, Vybrant Dye Cycle Ruby Stain (Thermo Fisher Scientific, Waltham, MA, USA) was added at a dilution of 1/100 and incubated for 30 minutes at room temperature. Fluorescence activated cell sorting (FACS) was performed using a FACSAria (BD Biosciences, San Jose, CA, USA) cell sorter. A live cell gate was selected based on forward/side scatter properties, and samples were gated IA6 (CD49f)-positive or –negative based on isotype control staining (RatIgG2A-isotype-control eFluor450, Ebioscience, San Diego, CA, USA). Analysis was performed using FACSDiva software (BD Biosciences, San Jose, CA, USA). Cells were sorted using high-pressure settings and a 70 μm nozzle and collected into filtered seawater.

### Quantitative RT PCR

Sorted cells were pelleted at 700g for 10min, and RNA was extracted using the Nucleospin RNA XS kit (Macherey Nagel, Bethlehem, PA, USA), which included a DNAse treatment step. RNA was reverse transcribed into cDNA using random primers (Life Technologies, Carlsbad, CA, USA) and Superscript IV Reverse Transcriptase (Life Technologies, Carlsbad, CA, USA). Quantitative RT-PCR (Q-PCR) was performed using a LightCycler 480 II (Roche Life Science, Penzberg, Germany) and LightCycler DNA Master SYBR Green I detection (Roche, Penzberg, Germany) according to the manufacturers instructions. The thermocycling profile was 5 min at 95, followed by 40 cycles of 95 °C for 10 sec, 60 °C for 10 sec. The specificity of each primer pair was determined by BLAST analysis (to human, *Ciona* and *Botryllus* genomes), by melting curve analysis and gel electrophoresis of the PCR product. Primer sequences are listed in the Supplemental file “primer sequences”. Relative gene expression analysis was performed using the 2^-ΔΔCT^ Method. The CT of the target gene was normalized to the CT of the reference gene *actin*:ΔC_T_ = C_T (target)_ – C_T (actin)_. To calculate the normalized expression ratio, the ΔC_T_ of the test sample (IA6-positive cells) was first normalized to the ΔC_T_ of the calibrator sample (IA6-negative cells): ΔΔC_T_=ΔC_T(IA6-positive)_ - ΔC_T(IA6-negative)_. Second, the expression ratio was calculated: 2^-ΔΔCT^ = Normalized expression ratio. The result obtained is the fold increase (or decrease) of the target gene in the test samples relative to IA6-negative cells. Each qPCR was performed at least three times on cells from independent sorting experiments gene was analyzed in duplicate in each run. The ΔC_T_ between the target gene and *actin* was first calculated for each replicate and then averaged across replicates. The average ΔC_T_ for each target gene was then used to calculate the ΔΔCT as described above. Data are expressed as averages of the normalized expression ratio (fold change). Standard deviations were calculated for each average normalized expression ratio (n=6).

### Mitomycin C treatment and rescue

Vessel fragments were cut and soaked in 60μM Mitomycin C (Tocris, Bristol, UK) in filtered seawater for 24 hours. MMC was removed and fragments were returned to flowing seawater for 24 hours before being micro-injected with cells (different numbers of IA6+, IA6-, G0 or G2M, as indicated) isolated from the blood of normal, healthy colonies. Rescue efficiency (number of fragments reaching stage 5) was scored after 20 days.

### Fluorescent *in situ* hybridization on cryosections

Vessel fragments at different stages of regeneration were fixed overnight in 4% paraformaldehyde in PBS at room temperature. Samples were washed in PBS and soaked in 15% and 30% sucrose for 30 minutes each before embedding in OCT medium. 20μm sections were cut using a Leica cryostat. Fluorescent *in situ* hybridization was adapted from ^(Langenbacher et al.)^ for cryosections. Briefly, *B. leachii* homologs of genes of interest were identified by tblastn searches of the *B. leachii* EST database (https://www.aniseed.cnrs.fr/aniseed/default/blast_search) using human or Ciona (when available) protein sequences. Primer pairs were designed to amplify a 500-800 bp fragment of each transcript (Primer sequences are listed in the Supplemental file “primer sequences”). PCR was performed with Hotstar DNA Polymerase (Qiagen Germantown, MD 20874) and products were cloned into the pGEM-T Easy vector (Promega, Madison, WI, A1360). *In vitro* transcription was performed with SP6 or T7 RNA polymerase (Roche, Penzberg, Germany 10810274001, 10881767001) using either digoxigenin, fluorescein or dinitrophenol labeling. Cryosections were air-dried and fixed with 4% PFA for 10 minutes and washed with PBS/1%Triton-X-100. Probes were diluted in hybridization buffer and hybridized at 65 C for 30 minutes. Probes were removed and slides washed with 2xSSC/1%Triton and 0.2xSSC/1%Triton for 15 minutes each at 65C. HRP-conjugated anti-digoxigenin antibody (Roche 1/1000), HRP-conjugated anti-fluorescein antibody (Roche 1/500) or unconjugated anti-DNP antibody (Vector labs 1/100) followed by anti-rabbit-HRP (Abcam, Cambridge, UK, 1/200) were used to detect labeled probes. Fluorophore deposition was performed by incubating slides for 20 minutes at RT in Tyramides (Alexa488-Tyramide, Alexa-555 Tyramide or Alexa-594-Tyramide, all from Thermo Fisher) diluted 1/100 in PBS with 0.001% hydrogen peroxide. Slides were washed twice with PBS and Nuclei were stained with Hoechst 33342 (Life Technologies). Imaging of labeled samples was performed using an Olympus FLV1000S Spectral Laser Scanning Confocal. Image processing and analysis was performed using FIJI. Quantification: images from 4 independent samples per time point were taken with a 20x objective and *ia6/h3* double positive cells, *ia6* single positive cells and *h3* positive cells were counted using the cell counter feature in FIJI. Counts were normalized to the number of nuclei for each image. Graphs represent cell counts in percent of nuclei, averaged over 3 independent samples per time point. Between 2500 and 3000 cells (Hoechst-positive nuclei) were counted for each time point.

### Tracking of EdU labeled cells

For every 10 zooids, 2μl of 1mM EdU (Thermo Fisher) dissolved in filtered seawater was injected into the blood stream of healthy colonies. After 24 hours, Integrin-alpha-6-positive cells were isolated from EdU-injected colonies by flow cytometry and injected into recipient vessel fragments 24 hours after Mitomycin treatment (see above). The experiment was performed two times. At stage 3 (n=10) and stage 4 (n=10) of regeneration, injected samples were fixed and cryosections were prepared as described above. Sections were fixed with 4% PFA for 30 minutes and treated with proteinase K for 7 minutes. Sections were post-fixed with 4%PFA for 20 minutes and blocked with PBS/1% Triton and 3% BSA for 1h. The clickit-reaction cocktail was prepared according to the manufacturer’s instructions and the reaction was stopped after 1h at room temperature. Nuclei were stained with Hoechst 33342. Regenerating samples injected with EdU were used as positive control, and uninjected regenerating samples were used as negative controls. Imaging of labeled samples was performed using an Olympus FLV1000S Spectral Laser Scanning Confocal.

### Small Molecule Inhibitor Treatment

Vessel fragments were cut and placed in the bottom of 24 well plates. 1ml of filtered seawater containing the Notch-signaling inhibitor DAPT (Tocris, 0.2-2μM), the Wnt-signaling inhibitor endo-IWR1 (Tocris, 0.1-1μM) or control drugs Exo-IWR (1uM) and the Notch-sparing gamma-secretase modulator E2012 (1uM) were added to filtered seawater and replaced every other day. The number of colonies that reached stage 4 were counted at 14 days post injury (n=16 for each condition).

### Pou-Phylogenetic analysis

Protein sequences used for phylogenetic analysis were downloaded from the NCBI database or determined in this study. POU protein sequences were aligned using the ClustalW algorithm in the MEGA7 application (Kumar et al., 2016). Phylogenetic analysis of POU family members was performed with the software RAxML using a maximum likelihood method, the JTT substitution matrix, and empirical frequencies (Stamatakis, 2014). RAxML software was accessed using the CIPRES Science Gateway (Creating the CIPRES science gateway for inference of large phylogenetic trees. In: Proceedings of the gateway computing environments workshop (GCE), New Orleans, LA, USA; 2010. p. 1–8.) and trees were visualized using the Interactive Tree of Life website (Letunic I and Bork P (2006) Bioinformatics 23(1):127-8 Interactive Tree Of Life (iTOL): an online tool for phylogenetic tree display and annotation).

## Competing interests

The authors state no competing financial interests.

## Acknowledgements

The authors would like to thank Bill Smith for providing facilities to grow *Botrylloides* colonies. We also thank Ben Lopez and the NRI-MCDB microscopy facility at UCSB. Shane Nourizadeh, Jessa Alcaide and Phillip Ahn are acknowledged for help with cloning. Delany Rodriguez and William Jeffery are acknowledged for helpful discussions and critical input. We would like to thank the reviewers from a previous submission for their constructive criticism and helpful input, which have greatly improved this manuscript.

## Notes

#### Summary of Updates

Changed presentation of in situ hybridization data and EdU-data. Edited text in introduction, results and discussion for clarity. Added table with raw data of cell transplantation experiments.

