## supplemental information for "Integrin-alpha-6+ Stem Cells (ISCs) are responsible for whole body regeneration in an invertebrate chordate"

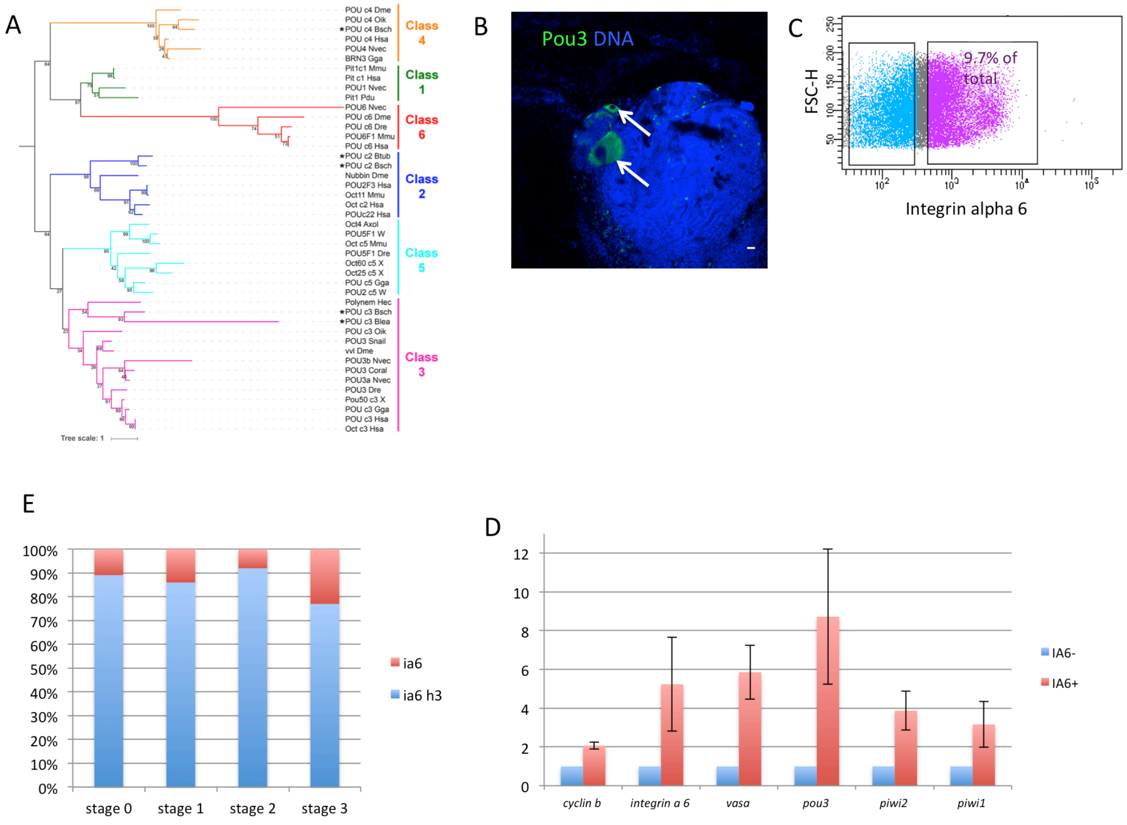


**Figure S1:** A: Maximum likelihood phylogenetic analysis of POU family members. Phylogenetic tree was constructed with RAxML using a maximum likelihood method, the JTT substitution matrix, and empirical frequencies. Botryllid POU proteins identified in this study are marked by asterisks. Families of POU proteins are labeled and indicated by vertical lines. Nodes are labeled with bootstrap values in units of percentage. The scale bar for branch lengths indicates the mean number of inferred substitutions per site. B: FISH for pou3 (green) on whole colonies. Pou3 is expressed in germ cells associated with the secondary bud (arrows). C: Fluorescence-activated cell sorting (FACS) of Integrin-alpha-6-positive cells. Integrin-alpha-6-negative cells were gated based on isotype-control staining. D: quantitative-real-time-PCR analysis of gene expression in FACS-sorted IA6+ cells. Data are expressed as averages from 4 experiments, normalized to IA6- cells. E: Percentages of ia6/h3 double positive cells (blue) and ia6 single positive cells (red) for each stage. Single positive (*ia6* or *h3*) and double positive cells were counted using the cell counter feature in FIJI, and for each stage, 4 images from 4 independent samples were counted.

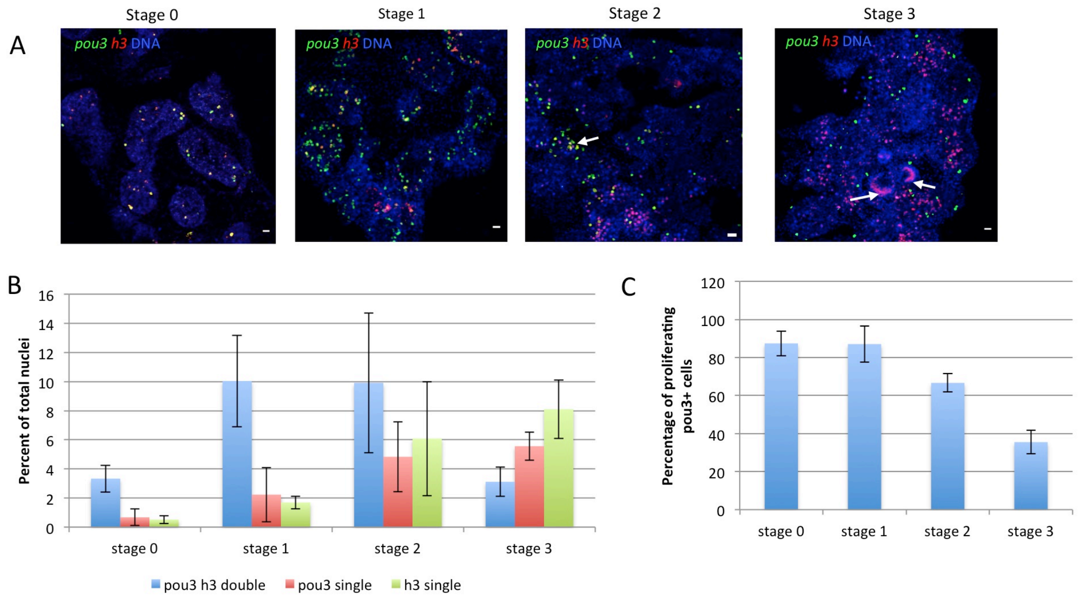


**Figure S2**:A) FISH showing expression of *pou3* (green) and *histone 3* (h3, red) during stages 0, 1, 2 and 3 of WBR. DNA was stained with Hoechst (blue). Scale bars 20um.Arrows in stage 3 indicate the beginning of double vesicle formation. B) Single positive (pou3 or h3) and double positive cells were counted using the cell counter feature in FIJI, and for each stage, 4 images from 4 independent samples were counted, comprising a total of 2500 - 3000 cells for each sample. Graph shows percentages of pou3/h3 double positive and pou3- or h3- single positive cells among all Hoechst-positive nuclei. Error bars show standard deviation. C) Averages of percentages of proliferating pou3+ cells among all pou3+ cells. Double positive cells were counted using the cell counter feature in FIJI, and for each stage, 4 images from 4 independent samples were counted, comprising a total of 2500 - 3000 cells for each sample. Error bars represent standard deviation.

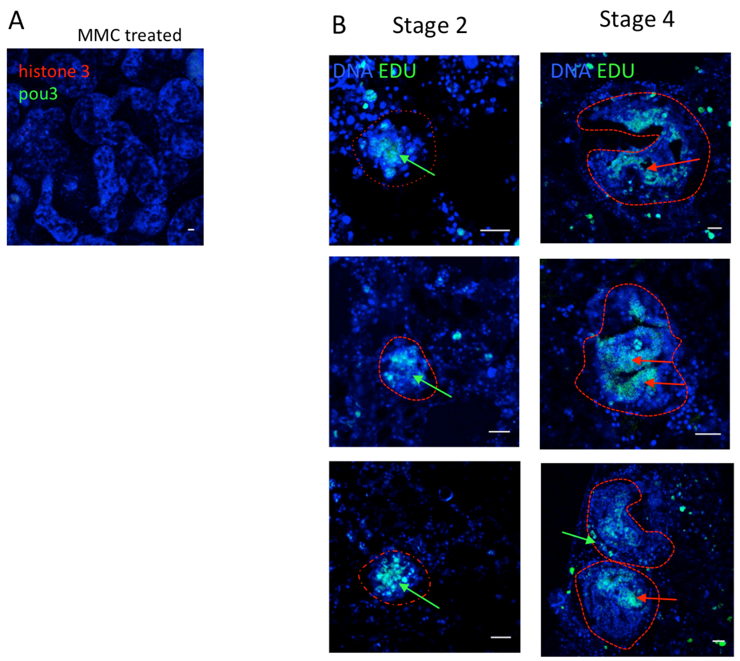


**Figure S3:** A: FISH for pou3 and histone 3 5 days after treatment with Mitomycin C. Scale bar 20µm. B: IA6+ cells were isolated from Edu-treated animals and injected into MMC treated vessel fragments. Overlay of nuclear staining (Hoechst, blue) and Edu (green) shows Edu-positive cells giving rise to regeneration foci at late stage 2 (white arrows). Red dotted lines outline regeneration foci. Scale bars 20µm. Images are representative of 10 samples from two independent experiments. Several samples from the same experiment as in A were followed to stage 4 (organogenesis). Red arrows: Edu-positive cells are present in several layers of differentiating tissue (red arrows) and occasionally in the outer epithelial internal tissue layers. Green arrows: Edu-positive epithelial layer. Red dotted lines outline regeneration niches. Scale bars 20µm.

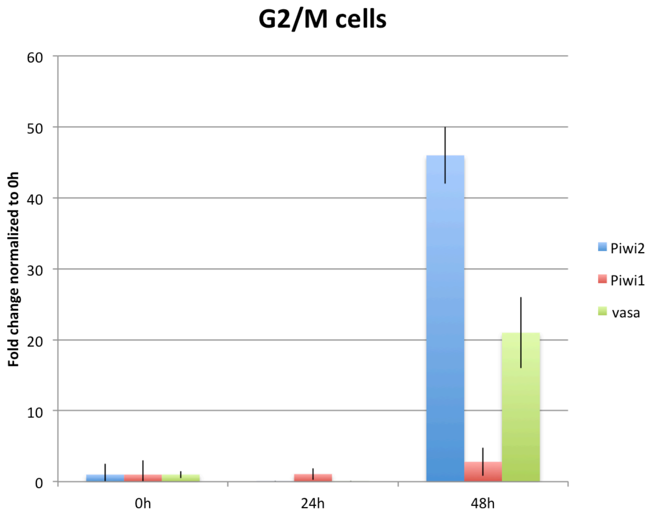

**Figure S4:** Q-PCR analysis of gene expression in cycling G2/M cells at different time points of WBR. Data are expressed as averages of 3 experiments, normalized to 0h. Error bars show standard deviation.

### Pou3-phylogenetic analysis – protein sequences

*Botryllus schlosseri* Class 4 POU (POU_c4_Bsch):
FESLTLSHNNMVALKPILTTWLELAEEEYRRKMEQSGLAEKKRKRTSIAAPEKRSLEAYFLVQPRPSSEKIAAI
AEKLDLKKNVVRVWFCNQRQKQKRMKFSAFNGENGGM

*Botryllus schlosseri* Class 2 POU (POU_c2_Bsch):
MLQYQQHRMFEDRRHSGGEFIHARPPSPGMHPGISQSPXHQSDYEETSSKRQRYEDEASELESLERFAKDFK
QRRIKMGFTQGDVGVAMGRFYGNDFSQTTISRFEALNLSVKNMGKLKPLLERWLIDVDRAISTGERSEGRPV
LSQPMMAMNSQSCAGRKRKKRTSIPTEGKSRLEDAFLKNPKPTTEEIGKFSEDLNMDREVVRVWFCNRRQK
QKRIATQQQRYVGEQHSPSSPAGFNEPHSVEGQRSPISSEDENYGSHHLSDRPPVHQAIEHSEMGTLPPQLPP
RPGIIQLKLSESHQMQFGRPPNYSSHSSLMGGNSAI

*Botryllus schlosseri* Class 3 POU (POU_c3_Bsch):
VEFYEMLSQVPNGELAYGSPLQDGCGKYSSHSEVGKCRNKTRIHENPHMPNHSPVMSYSTPGLSTYACLDSQ
PPIRSDTSQEQESFTKISDEHYRPYPNGYQFSNHCYQFNQSGYHRALPIPSLQEQLYSVHDSPRQRIPSRSPHTS
QVTEDFVIKESPAYTNDANVQSWHSFVHSAREISESSRLNTTSNAGCYPIANSVCASDKAGCVYSYQNQPGQY
PYCYSRNYYPASTNLQNRTWNPRPIDLSLKQDHCDGYGEMDYQPTHFTSYSPLRIDERNLLSDERTLNKLPC
DDMSLNGWTEEDMRQFSKVFKHRRTKLGYTQSDVGTSLGELYGSVFSQTTICRFEAQQLSLKNMCKLRPLLS
RWLQHKDNKHETLTPDIDQIDSENGPGRKRKKRTSIEAEVKAVLEKHFKLKPKPMTQEIVSIAEQLSLEKEVV
RIWFCNRRQKEKKVNEQVMRSQNPT

*Botryllus tuberatus* Class 2 POU (POU_c2_Btub):
LQGKEQKQKKLDYSQEDDRFAPAPGFHERHFVRQPSDFDVDPRQIRPDFAIRHSMLSHPVLLDNRRCSEEN
MSPPGPESPTVTPANMARHIPQSFSSDDQCDYDEIPKKRRMYDDEASELEDLEQFAKDFKQKRIKMGFTQGD
VGVAMGKFYGNDFSQTTISRFEALNLSVKNMCKLKPLLERWLIDVDRAISDRGEGGRLAISQPVIPMHPQQCT
GRKRKKRTSIPTEGKTRLEDAFKKNPKPTTEEICKFSENLKMDREVVRVWF

*Botrylloides leachii* Class 3 POU (POU_c3_Blea):
NNFYAHVSLTNENGRSVIAQGNECSGKQSPEYKEYETPIKELTYMSLHPRTVTSNGYYSQSSTFGENFRHFPT
QSPYTGHDHNPGYNYNSYPPTLIPADCLQGSLQSRHSFNSTFAPESYASEQNSDHPRISRSAPPNVTITECSDT
HYNANSVKCFSDRHSEYSYPHLPSPANMGVVCKREITTPSPQLRHEISSSWRGQELCPPSYLHQDTPQYRYSY
QANYWPLSPANSTSCSYQKASSNRFVKQERFQDYQTNTPMQKLPFRNFCTAAERGFSNETQFEQKFTSDES
MIGTESSDDMRIFANVFKARRIKLGFTQHDVGLDLKKFQGSAFSQTTICRFEAGGLSIKNMNRLKPLLTMWL
RHNDTEHISTLRDTDRSPLDNATTRKRKKRTCIEPQTKLALEEKFRNDQKPTTVQIAKIAEELSLDKEVVRIW
FCNRRQKEKKATVEIVQRDVA

### Pou-Phylogenetic analysis

Protein sequences used for phylogenetic analysis were downloaded from the NCBI database or determined in this study. POU protein sequences were aligned using the ClustalW algorithm in the MEGA7 application [1]. Phylogenetic analysis of POU family members was performed with the software RAxML using a maximum likelihood method, the JTT substitution matrix, and empirical frequencies [2]. RAxML software was accessed using the CIPRES Science Gateway (Creating the

CIPRES science gateway for inference of large phylogenetic trees. In: Proceedings of the gateway computing environments workshop (GCE), New Orleans, LA, USA; 2010. p. 1–8.) and trees were visualized using the Interactive Tree of Life website (Letunic I and Bork P (2006) Bioinformatics 23(1):127-8 Interactive Tree Of Life (iTOL): an online tool for phylogenetic tree display and annotation).


1. Kumar, S., G. Stecher, and K. Tamura, *MEGA7: Molecular Evolutionary Genetics Analysis Version 7.0 for Bigger Datasets.* Mol Biol Evol, 2016. **33**(7): p. 1870-4.
2. Stamatakis, A., *RAxML version 8: a tool for phylogenetic analysis and post-analysis of large phylogenies.* Bioinformatics, 2014. **30**(9): p. 1312-3.


**qPCR primers:**

actin Forward: AACAAGAAATGCAGACCGCC
actin Reverse: GGCCGATTCCATACCCAAGA
PCR product length: 146

cyclin b Forward: CCCCGCATGAGAACCATACTT
cyclin b Reverse: GCAAGCTGTAACTTGGTGCG
PCR product length: 146

Integrin-alpha-6 Forward: GGGATAGTAGTGCTGACGGC
Integrin-alpha-6 Reverse: CAGCGATGTCGGGATAACCA
PCR product length: 156

Pou3 Forward: CGATCTGTCGATTCGAGGCT
Pou3 Reverse:GGTTGCGTTGTCTAATGGCG
PCR product length: 146

Vasa Forward: ACTGTTGGATCTCAGCCGTG
Vasa Reverse:TCTGGCTCAAAGCCCATGTC
PCR product length: 153

Notch 1 Forward: GAACTCACTGGAAGGTCGCA
Notch1 Reverse: ACACTCCACCAACAGCGATT
PCR product length: 145

Notch2 Forward: TGCATGAGTAGCCTGGAACG
Notch2 Reverse: AGGATCTTGTACGCCATCAGC
PCR product length: 158

Hes1 Forward: ACTTGGATACGGGTGTGTGC
Hes1 Reverse: GTCAACGTGTTTCCGCCATC

PCR product length: 155

Frizzled 5/8 Forward: GCGGAACAAAACCTGCAAGA
Frizzled 5/8 Reverse: TTTCTTCCAAGGCAGGGCAA
PCR product length: 156

Dishevelled Forward: AACCTCGCTATCGCCACAAA
Dishevelled Reverse: TGATGCATGTTGCCATTCGT
PCR product length: 154

Beta catenin Forward: AAGATGACGTAGAGCGCACA
Beta catenin Reverse: GTGCTGCTGGGATTCTGAGT
PCR product length: 158

piwi1 Forward: ACCAGGAAGCGACTGACTTG
piwi1 Reverse:GGAGCTCGTTGTCCCAAGAA
PCR product length: 156

Piwi2 Forward: TAGGGCGTGGGACACTTTTC
Piwi2 Reverse: GAGTGCAGTTCCACTCGACA
PCR product length: 149

FISH probe cloning primers:

Integrin alpha 6 Forward: TGGCGTTGTAATCGTCGTGA
Integrin alpha 6 Reverse: TGAGCAGTGCATTTCCCGAT
PCR product length: 764

Vasa Forward: TGTCCCGTGACTGTCCTAGT
Vasa Reverse: TGCAAACTTGCGAGCCTCTA
PCR product length: 833

Histone 3 Forward: GCGGTCGTCTACACACTTCG
Histone 3Reverse: CACAAGGATTGGGTGGCTCT
PCR product length: 565

Notch1 Forward: AGGCACTTGCATTGACGGTA
Notch1 Reverse: CCTTCACACCTTGGTCCCTC
PCR product length: 792

Notch2 Forward: AAGTGCCTCCGTTCAAGCAA
Notch2 Reverse: CCCTGGTTACACCGGCATTA
PCR product length: 653

***B. leachii* Piwi mRNA sequences:**

*B. leachii piwi1*
GATCTGTTGTATTAAGGTTGCTGCAGAAGTATATGTAAAGTAAGCAGTTGTTTTATGGT
TGAATAATGGCTGAGCAAAGG
GCGAACCCAGGAAGAGCGAGAGGAAGATCTCGTGGACGAGGCCAACCCCTAGTACCTGAT
GCCTCTGCTCAACAACCTGC
TGTGGGGAGAAGAGGAGGATCAGATGCAGAGGTTGGAAGTAGCATGATTGGCTCAGGCA
GGGGAATGCAAAGGGGTTTGC
ATCATTCCACACAAGGACCAAGGGGTGATGCCGGCGTCCGAGAAATTACTAGAGGAGTT
GGCCGCATGACTTATTCTGGT
AACCGTCAAGTCATGCTTCTTGGAGCAAGGAGAATGTTGGACCCATCCATTTTTGATACT
GGCACAAGGCCAGATCACAT
CAAGGACAAAAGAGGAAAGGATGGAAAGCCTGTTCAGATCTTGACTAATTACTTTCCAC
TCGTGTCATCCAAAGCATGGC
GTTTATATCAATACAGGGTGGACTATGAGCCAGATGTCGACCATAAAGGGGCACGCCGG
GGAATGCTCAGGGATCATCTA
GATCTGATTGGCAATACTTATATGTTTGATGGATCAACTTTGTACACATTGAGAAGACT
GCCAGAAGAGGTCAACATCGT
TCACTCCAAAAGACTTTCCGACAATGCAACGATCCAATTGAAAATCGCCAAGACTGCTGA
GCTACCGCCAAACTCGCCAT
TGAGCATTCAGCTTTACAATTTGATATTTCGCAGTGTATTGAAGAAGCTTGGGTTGGAG
CAGGTCGGCCGACACTACTAT
GATAGCCAGGCCAAGTTTACCGTCAACTGTGACAATATGAGATTTGAGCTGTGGCCTGGT
GTAATCACAAGTATTCTACA
GTATGAGAAAGAGAGCGTCATGATGTGCACTGAAATTTCCCATAAGATGATGCGGGTAG
AAAGTGTTCTGACCATTATGA
AACAAAAATATGAGCAGTCGAGGCAGAGACGAACCAACTTCCATCAGGAATGCCAGCAG
TTCTTGCTCGGACAAATTGTA
CTAACACGGTACAATCACAAAACATATCGTGTCGATGGCATCGAGTGGAAGATGAATGT
CTCAATGAAGTTCAAAAAGGG
AGACGAGGAGATTTCCTATGTTGACTATTATAAAACGCAATACAACATTACGATTAAAG
AGATGGATCAACCTCTGTTGT
TGTCTCGTCCAAAGAAAAAGGAAGCTAGAGATGGACTGGAAGCGATTCACTTGGTGCCC
GAATTATGTACTGTGACTGGA
CTATCAGATGAGCAACGGGCCAATTTCTCTGTGATGAAAGCACTCGCTGAACATACCAAA
CAGGGACCTGCCAAACGTGT
GAACGCTCTGAAGAATTTTATTCAGAGAGTGTGCGGACACCAGGAAGCGACTGACTTGC
TTAACAAATGGGGGCTTCATT
TTGAAAGAGATTTGGTAGGAGCACCAGGTCGCGTTCTACCTCCAGAGCAGCTCATGTTTG
GAGAGCGTAAGTGCATCAAT
GGAGGTCAGTATGCTTCTTGGGACAACGAGCTCCGCAACTGCAAGCTACTGAAATGTGTG
GACCTCCGAAATTGGCATCT
CATCTGTCGCAGACAGGACCAGCAGATGGCGAACGGTCTGATTCAAAAGATGTGCCAAGT
TTCAAAAAAATATGGGTTTTG

ATATGTCGCGCCCTGTTTTGACTTGCATCGAACAGGACAGGACTGATGTGATTCTGAGCA
CCATTAAAGACATATGTGCC
AAGTCTAGCAACTGTCAATTGATACTTTGCCTGCTGTCCAGTGATCGAAAGGAACGCTAT
GATGCTATCAAGAAAGTGTG
CTGTGTGGATGTACCGATTCCCACTCAGGTGGTCAAGACCAAAACAATCTCGAAACCTCA
GAGACTGATGAGTATTGCAA
CAAAGATTGCTATACAAATCAACTGTAAGCTTGGAGGGGAAGCGTGGGCTGTGAATATC
CCGCTTGGTGGTACAATGGTC
ATTGGAATCGACACTTACCATGACTCTATCCATAAAAGTCAGAGCGTCGGTGGGTTTGTT
GCCAGCATCAATCGAGGCTT
CACCAGATGGTACTCATCCACTACAACCCAGCAAGCAGGTGTTGAACTCATTGACGGATT
AAAAGTCTGCATGGTCGGGG
CCCTGAAAAAATACAGAAGTGAGAATGGTGAATATCCGAAAAAAGTTTTCGTATTCCGC
GATGGTGTTGGTGATGGCCAG
TTGTCCATGGTGAGAGAACATGAAGTGCCTCAAGTCATCCAATGCTTGAGAGACCCGTCC
AATCCGAATGCCAAACCTAT
TCCACTGTCTTATATTGTGGTGAAGAAGCGCATTAACACGAGACTTTTCACAAGTGGTA
GACAAGGCATGGCAAACCCAC
CTCCTGGCACCATTGTTGATGATGTTATAACTCGTCCGGAATCGTATGATTTCTTTGTGG
TGAGCCAGAATGTGCGGGAA
GGGACTGTGTCGCCTACACACTACAATGTAATATTTGATGAGTCCGGATTGGCTCCAAAT
CACATGCAACGCTTGGCATA
CAAACTCTGTCATGTCTACTACAATTGGCCGGGCACAGTGAGAGTTCCTGCTCCATGTCT
GTATGCTCACAAGTTGGCTT
TCCTTGTTGGTCAAAGCGTCCATCGAAATCCTGCAGCATCGCTGGCTGACAAATTGTATT
TCTTGTAATCATGTATCAAG
TGTGCTTTTCTCATAACGGTGTGCCTTCGTTGATAGTTCCAGTTTTTATCATTTTTGACA
CAAGGTGATTTGATCATGTA
TAGTGATAGCAGCTATACAACATTCTGTTATTTCTTGAGGTTTTGAACTGTTCCTGTTTG
TGCACTGCCTTTTTCTTGAG
CTGGGCGCGACCAGCCTGTAATTCGCCGATTGTGGGCGATGCCGTGTTGTGGTGATGATT
TCTTCGCTTGAAACTGGCCG
CTTGGTCAACTAGGGGTGCGTTTTATATTCATAAATGTGAAATTGATAATTGGATTGGA
TCTGTTACGAGCTAACGATAA
ACTGCCCGATTCATGCTGAACTGTTACATTCGCCAGTATGTGGTGATGATTTCTTCGCTT
GAAACTGGCCGCTTGGTCAA
CTAGGGGTGCGTTTTATATTCATAAATGTGAAATTGATAATTGGATTGGATCTGTTACG
AGCTGACGATAACTGCCCCTG
ATTCAGGCCGAACTGTTACATTCGCCAGTATGTGTGTAATCGTTTTATGTAACTTGGCGA
TTGTAACGTTTTATGATATG
CTGTGTTGGTTGGATTTCGTTGGTGAAATGAAGCTTGTCTGTCATGATTGCGTTGTTGCT
AGCTGGTCTTATTTGATGAC
AAGCGGGTGCACCTATTTTAGTTGGTGTTTGGAGTTTACGGGACTTCGGAAAAATTTGT
CCCTGATCACGAGGTTGATAC
TTTACCTCGTGAATTATTGCAATGCGATACGAATCCATTCTAGAACCGCCGCTGGTTGCG
GATGGTCACGGATTCCCTTT

CAGTTCATAATAATTTTTTGTTCAAGTAGGCCTATCGATTCAGCGTCGACTTATTAGCAT
GAAGCTAGTAGCATTAGCAT
GCCACAACAATCGTAGTCCACGATTTAGTCGGCGAAACGTGTCCATCCATAATCCTAATG
CAATAACGATCTGTCATCAG
GTGCACAAGCCCCATTTCAAGGATCCGGATATCAAATCCGGGCGGCCCTTGAGAGAGGTG
GCCTGCTGAACATTGGATTG
AGTTGCTTTATTAGAAATGTAAAAGCCTACACTACGTACGACCTAGACCCTGGCGTTCGC
TTCTCACGGGAAGTGAGAAA
ACCCGCCCCGAACTAGCTTCCGTTTCTCATTCACTCTGGTTTTCGATATGGGTATTAGAT
GCTAGGCCTATTACTATTGG
TCTGCAAACAGCCTACATCTAGACTAGCTCTGTCGCTTGCTTTGCCAGTTTTAATGCAAC
AAGCTCGATTTCTTCTTTGT
AAATGTTAGGACTGGTTCAGTGGTGCGAATTCACTCTTTTGGCTATTTTCAGGGCATTCT
GTGGAACCTAAACTAAGCTC
TGTTGCTACTATAAGCTAGGCTAGGACCGTGTATAGTCAGCTAGAGAGGGC

*B. leachii piwi 2:*
CCGAGATTACTGCAATCTCTGCCGTTGCAGGGTCCCTTGACACTCCGTCGATGTTTTCTC
TGAATATAAGCCAAATTCTC
TCTCAATAAAAAAATGTATCGCGAATTTTTTTGATTTTACAATCTCTTCGCTGCTGCCTT
ATTGTTTCATTTTGCATGGC
TTCCTCTGTCCCAGCCAATGTCGGTCGATAAAGCAGACCTATCTCATACAGGTACCGCGC
CTACAGTTCAACACCACACG
CCTAATCAGAGAAAACCCCTATGTTATATGACCCAATGATATATATGACAGATTAGAGA
GTCTGTAGGCCTACTGCCAGA
CCAGTGGCCGATGTTAGGCCACCTTTTCGAACTAGAACGGCGCAACTCCGTTGCCTATCC
ATGCGGAAAGGACTGAAGCA
CTGAGCCGAATCGTCCGGATCTTTGACACGGGGCTCTACACAAGATGGCGTGCGCGTACC
GTAGACAAAGACCTAACTGT
TCCAGGTCCCACGTGATGGACCCACCTGGTAGTGAACCCTGAACCGTTTGTTGGTTTGTT
GACTTGTTGAGCACAACGGA
GAAAAGGAAGTGGTACAGTAACTGGAAGGTGAATTAGTGAATATTTCGTTTATTATCGC
CATTTTGTGCTTTGAGGCGGA
GTTTATCCCGCGGACTATCCATCGCATCCGCATCCCGTTGAGGGCAAGCCCTCATTTCCT
TGATGCGAATGCATAGCTGT
TTTTCTATGTTGCTAGTAGCCTACGGGTGGAAGTTGTTGGTCGTTTGCAATACTGCTTGC
CCATATGTAAAAGTAGTAGG
CTATGATTAAAATGCCACCCACTACCAAAACATCTAAGCTGGCTGAGAATTATGCAATCT
AGTCTACCCGGTGTGCTAGT
CATGTTCGCCGAGAATCCGACCACCGTTTTGATGGGATAGATGCACTTGATGCAAAAACT
ACCTCAAGTCAGTCCACGCA
ATGTACGGATACATTTTGAGATCAATGTTTCAATTGAGTATCCGCACCCTAACACGGTCT
AGCCTTGGCTTGTAGAGTTT
ACCAAGTTACAAAAACGTCGCCTTTGTCGGCCTGGAGCCCAAAAATGTAGCAATAGGCTA
CAAAATGTACTGCTTATCCT

TGTGTTGAGAGCCTTGAAAGTACTGAAATGCGGAGGATAAGTTATGACGTCATATTGAT
ACATTTTCAAGTGTTACCAAC
AATGATATCTAGCATCTGCGCACCAATTTTGAGACCAATCAGGTCGGCCCGGTAGAACTC
AAGTTATAACATAGAAAATC
ATCTTCCAACTACATATTATAGATTTGCTGTGTTATGAGTATGAATCTCCACGCTCTGGC
AATTGTGGCAGCGCCTGGCC
GCCTGGCTGGGAAGCTACAGTCATCATATTTAACGCACTTAGTAGCATGGCAGCAAGTGC
CTCAACCAAATTAAAATACC
AATACAATGTTCCGGTGAAACTCTTCCAAAGGAATTTTATGTTTGATTTTTGCAAGGCA
TGCCTTTAAGCCGTGTCCGGA
TATGTGCGAGTAAGACTGAGCCTGGTTCATCTCATGTAAATACTTGAAAACAGATTTTA
TGCTTAGTAGCTGTCTGCTGC
CTTTGAATTGTGTATTTACATCACTTTTGCCTCATTGTACAAGGTTGCAATAGGAATTTG
TTATGGAAAGCCATGGGCTG
GGAAGAGGAGGCCGTGGGTATAATGCATTTGGATTGAGTAGTGGCGGCAAAGCCGGTCA
ACCCAATGAAAGTGAAGAGTC
ACAAGCAGGAAGAGGTGGTTTGCTGTTTCCAGGGCGTGGGAGCATTGGAAGAGGGGCAT
CTTTACTTCGGGGAATGAGTC
GACCAGGTCCATCTGTAGGGCGTGGGACACTTTTCCAAAAATTACTGGGGCCTTCTGCTG
GAGACAAAAGTGAAAGCTCA
TCAAGAATAAAAGCTGATGTGTCTGATGGCCGTAGCAGTAACACAGAGTCAATTTCTGA
AGATTTGTCGAGTGGAACTGC
ACTCACATCAAATGAGTCATCAAGTCGTAATCGAGCCAAAACAGATTCAGGATTTCATC
AAACTCACTCGAGTGGAGGAA
TCACTGGTGCCAGCGGTTCCGTTCAAGTAAGACCACCACCACCAGCATTCCGACTCATGC
AGACAGCATTCAGTTCAGTC
TCGCTTGGCAGAGGTTTCAACGTGCCCAGCTTAGGAAGAGGTCTTCGTATGTCTGGTGCA
AAAGCACCTATTGGAGTCAC
TACATCAGTTGGTCCAAAATCGACGGGCACGTTGTTTGAAGGTGGTGTTCAGACTCCAAG
ACAAACAGAAACTGGTCGGT
TTGAGGGTCCAAAAGAGAGTGAAACAGTTGTCCCAAGTCCACAGAGCTCTAAATCAGGA
GAGACTGTTGAACCTATCCAA
ACCCTGCCTGCATCTAAAACTAGGTCCCCAGAAATGAGGCAGCCGAGAATGGGTTTGTTG
CCTGGGAGGAATATGTCTTC
TGGTAGAGGTGTGACACTCCCTCATCACAAATTAGAACCGACTTTGGGAAGTGCATCAAT
AAGAAGTCTATCTACTTCAG
AAGCATCGGCTACTCATGAAAGGCCAATGAAACATGATCCACCTGTTATGATGGAAAGG
GCAGTCACTGAGTATCATGGC
AATAGTGGTGTGGAAATCCCAACACGTGTCAACTGTGTTCGTGTAAACTGCAGGAACCCT
GTCATTTATCAATATCATGT
CACTTTCAAACCAGATGTTGATAGTAGACAACTTCGGTTTTCCTTGATCAACCAACACAG
GGAGTTCATAGGACAGAAAG
CGTTTGATGGTGCATTGCTGTATCTAACTAAAGATATTGGTCAAGACACAACGCTGATT
TCAGAAAGAATTACAGATGGT
GCCAAAGTTAGCTTGAGAATTCAAATGACAGCGAAACTGAACCCGGCATCACATAAGTT
GATTCCATTTTACAACGTGGT

AATGAATAAGGTCATGAAAATACTCAAACTTTGCCCAGTTGGACGAAACTATTATGATC
CAGCATGCCCCAAGTCTGTTG
AGAAATACAGGCTTGACTTATGGCCTGGATATGTCACTGCTGTGAGGGAAATGGAAGGC
GGCCTATTCTTGAATATAGAC
TGCTCTTTTAAGGTTCTTCGAAATGAAACAGTACTGCAACAGATGCAAGAATTGTATAG
AACTTGCAAAGCTGACTTTCG
CGCAAGTTGCATTAAGGAATTGGTTGGAGTGACAGTGCTGACACGCTACAACAATAATA
ACTATCGCATTGATGATATTG
ACTGGTCGCTTTCTCCAATGAGCACATTCACAACCTCAAAAGGTGAAACCATCAGTTTCT
TTGACTACTATAAAAATCAA
TATGGAATTGAAATAAAGGATCTTAAGCAGCCAATGCTGATCAACCGGCCTAAAAAGGC
GACTCAGAGACATACCAGAGC
TGAGAAGGGTGAGGAGCGTGTGCTTTCTCTAGTACCGGAGTTATGCTGTTTGACGGGAA
TTAGTGATAAGCTCGCTGGAG
ATTTTTCAGCCATGAAAGACATTGGAGCAATCACTAGAATGGGTCCTGTTGACAGAAAT
CTGCATGTTGAGCAGTTTTTG
AAAAGAGTCAATTCATCTACTGAAGCAAGAGGAGAGCTGTTGAAATGGGGATTAGAACT
TGAATCCACTTGTGCTGCTGT
CAAGTCTCGGGTTCTACCTGGAGAGAAAGTCATGGTTGGTGGAAACAAGGTATTCCAGT
GCAATGAGCAAGTGGACTTTT
TCCGTGACATATCCCGTGCTCCAATGTTGAGATCGGTCCCATTAAAGAATTGGATAATCG
TGTTCACTTCTAGAGATCAG
TCCAAGGCAAGAGATTTTGCAACCAAATTTAATGAAGTCTCGAGGTGTATTGGAATGCA
AGTACATCCACCTGAGTTCAT
TGAACTGAAAAGCGATCGTAGTGATGAATTCAAGCACGCAATTTCAGGAAACATAAACA
ATAACCTTCAACTGGTTGTTT
GTATCTTTCCTACTCTTCGTGAAGATCGGTACAATGCGGTAAAAAGACTGTGTTGTGTTG
AAAAGCCTGTACCCTCGCAA
GTCATAATATCTAAAACTTTGCCGAAAGATGCCAAGATGGGCAAGTTTCGTTCTGTAAC
GATGAAAATCGCTCTTCAAGT
CAACTGTAAGTTAGGTGGTGAGCTTTGGGCTGTCTCAATTCCACTTGATGGACTCATGGT
TTGCGGGATTGATGTCTACC
ATGAAGGGAAAGGAGGCAAAGGAAGATCTGTGGCTGCGTTTGTTGCGAGCATGAATAAG
TTAGTTACAAGATGGCACTCT
CGAGCAATACAGCAACAATTGCCCAATCAAGAAATCATTGATGGCTTGAAGCAATGCTT
TTGTGATGCATTAACAAAGTA
TTACGAGTCAAACCATAAGTTACCTGATCGTATCGTAATCTATCGTGATGGAGTTGGAG
ATGGTCAATTGTCTGTCGTGG
CAGAGCATGAGGTACAGCAATTGAAGGATGCCGTGAAAATGTTCAATGAAATAGCAGGT
GGTGTATACGCACCTAAATTT
ACATTTGTGGTGGTTTCAAAGAGGATTAATATTCGTTATTTCAAGAGAGATGGAAATGG
TCTTGTCAACCCCCCACCAGG
AGCGGTGTTGGATCATACCGCTACGAGACCAAATTGGCCAGACTTCTATCTGGTGTCTCA
GCATGTTCGCCAGGGTACTG
TTTCACCAACGCATTATATTGTTGTTGACGGTTCCGAGAATATGAAACCTGATCACTTGC
AACGGCTTACCTACAAGCTC

ACCCATATGTATTATAACTGGCCTGGTACAGTCAGAGTGCCTGCACCATGTCAGTATGCT
CACAAATTGGCTTATCTCAT
TGGTGAGAATATACGAATGGAACCCGCTCAGGATTTGTGCGACCGTTTGTTTTACCTATA
AACGTTTTCACTGGTCCACA
GAGATGGTATAAACCTAATTATGAGGTATTTTTAGAATTTGAAATTTTTGACAAATTCC
TACAGGTTGCACTGGATAGTT
TTATAATTGCACTCCCTTGACATTCAGTCCGTGGAAGGCTGAGTTTGGATTCACAGCTCG
ACCCTCCCCGAGTGGGGGA
TGTGTCGTAGGAAGAGCGTCATGGTGCAAAACTACCCCTCAAACCTCATCTTGCTGAACT
GGATCAAAGCATCCCCTGTC
CACTACTTTCCCCCTTTTTACAAGCATTTTTTATTTTTTAATCAAGACATTTTTGATCAT
TGCAAATGACGCCATCTGTA
AAGTGGACCTGTTCCTTTGATTTGCACACGCTTCAATGTGTGCATGAGCTATGTTATCAT
TACTTTGATTATGATGGTTT
TGTCCTAAAGCTGTGTTTGCAGTTCCATTTATCTCATAGCCTTGTGGATTCAGTTCCGAT
GACAATGGTCGGAAATCATA
GTTCCACTTTGGAAAGTTTTACTTTATTTGGTGAGTCAAATTCGTTATGACTTTAGAAA
ATTGTGCATGATTGAGTTTCG
TTTCAAACTGCTCCCTGTTATGCTGCATAGTCTGGATTTTGTGCTCTTTTAGAATGTGTA
TTAATGTCAAAATCTGGACA
AAAAGTTGAGCAACCTTTGGCAATCAGTTAACTACCTGTGTATGACAGGATCGGAAATA
TAGGACGTATATTTATTCTGT
CCTTCAATTTCGCTGCATATACACAGATAATGATGATAGGCCTACAGACGACATGTGCTG
CGAATTTCGTTCCTTTTGTG
AGTTGAATACACGCACAAACAGCGGCTATTTGGGACGTGATCCGTAGGACAAAACTATC
TTTCCCTTTGGCCTATCTGTA
TCGGTGTGCGATATAGCATAACAGCCGAGGCGAGTTTGGTATGTTTCTGCGTCTGATGGT
TATCAGCAGGGACTAATGTT
GATGGTGGTGGGGTCTTAAGACAGAAAGACACCAAACTCTGTGAAGTTTGGATGAGGCT
TGTATGGCGGAAGACATGAGG
TTGTACTTTGAGTTTCCCACTGACTGAGGCACTGACCTTATGCTAAAGGTCTAGTCGGCA
CCGCGAGTATGTTGCTGATG
CTGTCAAGCAATTCCAGGCCATCGCACGACGCGGCAGACATAAAACCCACAAAGCCGAGG
CAGTGAGTAGGCCTTCGGTG
TGTACTTAGTGAGTGCGACCATGATAGGTACTATGACTTGCGGCGTTTCACACAGCTCGA
TAAAAGTGGAGCGATAATTG
ACCGTCATTGTCGATTCCTTGATCGATGATTCAAAAAGATGGAAGAAGCCGTTTTTTAT
GTCAAGACAAACACGAACAGC
TTTTGCGAGATATGTCTTGCCATTATGCATTCATTAGTCTTGGTATTGTATAGTAGACCT
TTGAAGCAGATATGGTTTCC
GGATGAGTATTTCTGGTTTTTACACATCAACGGCTTACGCCAAGTCACCAGCCCAAGCTT
CAACAGAATAACGCAGGCAG
ATTGGCCCCTGTCGTGCTAATTTTTCACGTGTTTTTCCAGGAGTTGTGGCCACAGATGCC
ATAGTCATACAATACAGTTG
TCTAAGACTTTACAGCAGACACTCCTGCGGGAACATGCCGCATCCTGCCAGATGCGATGT
ACTGATGCGCATCACGCAAT

CTTTGCACGTTGCAACTCCTCATGACTGGTGAAAGCTTGCCGCTCGTTCATTACAATGAA
GGTAAAATATACCGCTTTTA
GATTCAGATGTATGCATGTGTACATTTTCTATGTACCTTCCCTTCTCTATAGGCCTGATA
ACAGCGCCGAGATATCCATC
CTCTCTGACCACTGAGCCAGGTAACAACTCAAGCAACACGAACTCATGTCGTCACGGGAA
CTTCGTGTTCGTTCGAGGTC
AAGCTGACGCTGAGTACCAGTACCGTACTGTTCTCGAAAATGTATGCAGCGGGATTGACG
TGACGTCTTTCCGTGAGGTG
TCGCCTAGGGATTGCGCGGTGATACGGTAATCAGTCTAATCGTAGAACGTAATCGATGA
TATACTTCGCTTACGGTACAT
TGCGGCCGTTTTGGCAGCCAGTGGTAACAGGCACACGGTTCTTGTCGCCTAATGTGTAAG
AGAGTACCGGTACAACGTAT
TTCCGTTCGAATGTTTCGCACCAACTTATGAACGAGATTTCACTTTTAGACGTGTAGCCT
AGCCTACAACTTGAAATGCT
CTGTTTCCGCGAATCTTCGTAACTTCCATTTTCGGGGATATCACAACAGAAGCCGAGCTC
AAGCAACTAGAAACTACCCA
ATAGGATGAGTCTCGTAAGTTTTTTTATCTGTTCAACGGGTTCCACCGATACAGTACTT
CACGACAGTTGCGAGTAGTG
TTTCAGCGAGTCACCATCTGCGCTGTAGGCCCAAGTGGTTATTAGGCTAACGCTAAGCCT
ATATCTAGTCTCACCGAGTT
GTTCGAAAATCGTGGCAATCGTATCTTTGAAATTTTATATCGTCAAATGGGAATCACGA
AGCACTTTTCGTAACGATACA
TATCTATGGCAAATTTGACAGCTCATGGGGCATAGGCTATCAGAGCGATCGCTTGTGAA
AGAAGCAGATACTCACCGCGT
CGGCTTGGTGTAGATACAGGACCGTGATAACAGCGACGAAAAGTAGTTTTGTGT
